# Investigation of the validity of two Bayesian ancestral state reconstruction models for estimating *Salmonella* transmission during outbreaks

**DOI:** 10.1101/574087

**Authors:** Samuel J. Bloomfield, Timothy G. Vaughan, Jackie Benschop, Jonathan C. Marshall, David T. S. Hayman, Patrick J. Biggs, Philip E. Carter, Nigel P. French

## Abstract

Ancestral state reconstruction models use genetic data to characterize a group of organisms’ common ancestor. These models have been applied to salmonellosis outbreaks to estimate the number of transmissions between different animal species that share similar geographical locations, with animal host as the state. However, as far as we are aware, no studies have validated these models for outbreak analysis. In this study, salmonellosis outbreaks were simulated using a stochastic Susceptible-Infected-Recovered model, and the host population and transmission parameters of these simulated outbreaks were estimated using Bayesian ancestral state reconstruction models (discrete trait analysis (DTA) and structured coalescent (SC)). These models were unable to accurately estimate the number of transmissions between the host populations or the amount of time spent in each host population. The DTA model was inaccurate because it assumed the number of isolates sampled from each host population was proportional to the number of individuals infected within each host population. The SC model was inaccurate possibly because it assumed that each host population’s effective population size was constant over the course of the simulated outbreaks. This study highlights the need for phylodynamic models that can take into consideration factors that influence the characteristics and behavior of outbreaks, e.g. changing effective population sizes, variation in infectious periods, intra-population transmissions, and disproportionate sampling of infected individuals.

## Introduction

Ancestral state reconstruction models estimate the ancestral states of organisms based on their evolutionary history. Outbreaks are “…the occurrence of disease in excess of what would normally be expected in a defined community, geographical area or season” (1). Ancestral state reconstruction models have been used to investigate the transmission of infectious agents between animal populations over the course of outbreaks, with host population as the state (2). However, as far as we are aware, no studies have validated these models for this type of analysis.

The discrete trait analysis (DTA) and structured coalescent (SC) models are ancestral state reconstruction models. Both models treat each host population as a discrete trait and can be approximated using Markov chain Monte Carlo methods (3,4). There are many differences between these two ancestral state reconstruction models. In the context of host association studies, the DTA model uses a substitution model to model the transmission between host populations (3). The pruning algorithm (5), often used in phylogenetic analysis to account for possible mutations, is similarly used by the DTA model to integrate all possible migration histories (6). The SC model assumes that the pathogen associated with each host population has a fixed effective population size and models the transmission between populations. The DTA model assumes that the number of offspring an individual pathogen is likely to produce is independent of its host population, whilst the SC model allows for variation between host populations (4). The DTA model assumes that the proportion of isolates sampled from each host population is proportional to the size of the pathogen population associated with that host, whilst the SC model allows for variation in these population sizes (6). Some of these assumptions are applicable to the investigation of outbreaks (e.g. varying effective population size), whilst others are not (e.g. isolate proportionality).

Salmonellosis is an intestinal infection caused by non-typhoidal *Salmonella* strains. Salmonellosis outbreaks vary in size and can involve one or more host populations (7). Identifying the amount of time *Salmonella* spends in a host population over an outbreak and the amount of transmission between host populations can inform control measures to limit salmonellosis outbreaks, e.g. if human cases are primarily from exposure to poultry sources then control measures that limit human exposure to poultry or decrease the amount of *Salmonella* in poultry may be beneficial. However, there is growing evidence that exposure to human sources contributes more to salmonellosis outbreaks than previously thought (8). Therefore, methods and models are required that can approximate the number of cases that are the result of exposure to different animal and/or human sources. The aim of this study was to use simulated outbreaks to investigate whether the DTA or SC models could be applied to infer transmission dynamics in outbreaks involving multiple hosts, motivated by non-typhoidal *Salmonella*.

## Methods

### Outbreak simulations

The MASTER package (9) in BEAST2 (10) was used to simulate stochastic transmission dynamics for a pathogen infecting structured populations, including associated phylogenetic and transmission trees. Outbreaks were generated using a stochastic Susceptible-Infected-Recovered (SIR) model, intended to simulate the transmission of zoonotic salmonellosis. In this model, susceptible host individuals become infectious by exposure to other infected individuals:

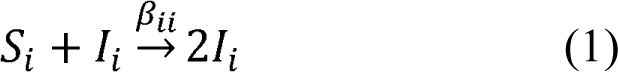

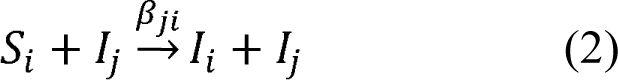

Equation 1 represents the transmission of the infectious agent from an infected individual to a susceptible individual of the same host population. Equation 2 represents the transmission of the infectious agent from an infected individual to a susceptible individual of another host population. Here, *S_i_* represents a susceptible individual from one host population, *I_j_* represents an infectious individual from the same host population, *I_i_* represents an infectious individual from another host population, and *β_ii_* and *β_ji_* represents the transmission rate per susceptible individual per infectious individual.

In this model, infectious individuals also recover or are removed over time:

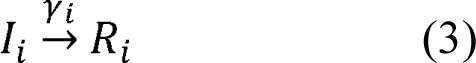

Equation 3 determines the infectious period for an infectious individual. Here, *I_i_* represents an infectious individual in one host population, *R_i_* represents a recovered/removed individual in the same host population, and *γ_i_* represents the recovery/removal rate per infectious individual for this host population. The mean infectious period for a host of type *i* is 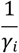.

### Simulated outbreaks

We simulated 23 outbreaks using the MASTER package, hereinafter ‘outbreak simulations’. This created 23 transmission trees consisting of all the transmissions that took place over the course of each simulated outbreak (Fig 1). These simulations consisted of two host populations: human and animal. We wanted to compare the simulated outbreaks with a previously reported salmonellosis outbreak in New Zealand that involved *Salmonella enterica* serovar Typhimurium DT160 (herein, DT160) (11). Therefore, the initial susceptible host population size, infectious period (*γ*) and transmission rate (*β*) values varied between the 23 simulations but represented possible values for salmonellosis outbreaks in New Zealand (S1 Appendix).

**Fig 1.**
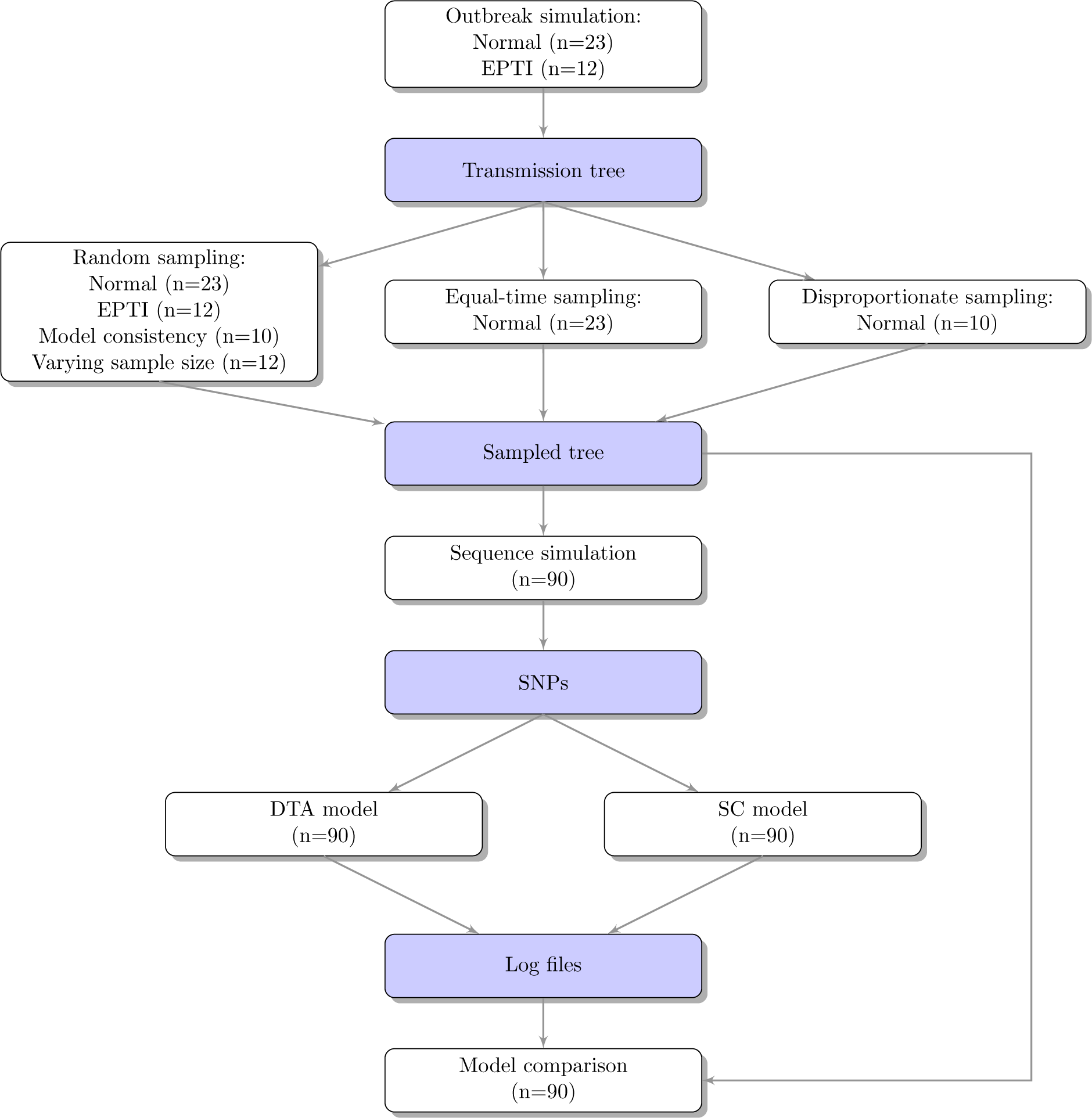
Flow diagram of the methods used to compare the SC and DTA models using various sampling methods. White rectangles represent the methods used and blue rectangles represent the data produced.

### Simulated genetic sequences from outbreaks

One hundred ‘*Salmonella*’ isolates were randomly sampled from each outbreak simulation, after stratifying for host population, hereinafter ‘random sampling’. For each outbreak simulation, the transmission tree was simplified to only include nodes common to the 100 isolates (both steps were accomplished using custom Perl scripts). The sampled transmission trees were used to simulate genetic data for the 23 simulated outbreaks using the sequence simulation capability of the BEAST 2 package MASTER, hereinafter ‘sequence simulations’. 800 SNPs were simulated in total for the 100 isolates, similar to the 793 core SNPs shared by 109 DT160 isolates (11). Perl and R scripts were used to analyze the sampled transmission tree and to calculate the amount of time spent in each host population and quantify the number of transmissions, later referred to as the ‘known parameters’.

### Model consistency

To investigate variation in model estimates between different samples (i.e. model consistency), one of the simulated outbreaks was randomly sampled 10 times after stratifying for host population. For each sample, sequence simulations were used to create genetic data.

### Sample size

To investigate the effect of different sample sizes on the models’ estimates, one of the simulated outbreaks was randomly sampled 12 times. The number of isolates sampled systematically ranged from 25 to 300 isolates in 11 increments of 25. For each sample, sequence simulations were used to create genetic data. The genetic data systematically ranged from 200 to 2400 SNPs in 11 increments of 200, respectively. To determine if sample size affected the extremity of a model’s estimates, the simulated outbreak chosen had significantly different population values between host populations and similar transmission values for comparison.

### Disproportionate sampling

To investigate the effect of the relative number of isolates from each source on model estimates (i.e. disproportionate sampling), as expected during the outbreaks, one of the simulated outbreaks was randomly sampled 10 times with different numbers of animal and human isolates. For each sample, 100 isolates were analyzed, but the proportion of isolates that were from each host population were systematically ranged from 5-95% in 10% intervals. For each sample, sequence simulations were used to create genetic data.

### Equal-time sampling

To investigate an alternative sampling method, ‘equal-time sampling’, an in-house Perl script was used to stratify the isolates from the initial 23 simulated outbreaks by host population, before randomly sampling an equal number of isolates from each year of the simulated outbreaks, to a total of 100 isolates. Sequence simulations were used to create genetic data for the samples.

### Equal intra-population transmission and infectious periods

To investigate if different intra-population transmission rates and infectious periods had any effect on model estimates, twelve additional outbreaks were simulated but with equal intra-population transmission rates and infectious periods (EPTI) for both host populations, but inter-population transmission rates and initial susceptible host population sizes that varied. For each simulation, 100 isolates were sampled using random sampling, and sequence simulations were used to create genetic data.

### DTA model

For the DTA model, the genetic data was imported into BEAUti 1.8.3 to create an XML file for BEAST 1.8.3 (12). The generalized time reversible (GTR) model was used to model base substitutions (13), the Gaussian Markov random field (GMRF) Bayesian skyride model was used to allow for changes in the effective population size (14), and a strict molecular clock was used to estimate the mutation rate, which was calibrated by the tip date. The XML file was run in BEAST for 10 million steps as a single run with a 10% burn-in.

### SC model

For the SC model, the genetic data was imported into BEAUti 2.4 with the MultiTypeTree package (4) to create an XML file for BEAST 2.4 (10). The GTR model was used to model base substitution and a strict molecular clock was used to estimate the mutation rate, which was calibrated by the tip date. The XML file was run in BEAST for 250 million steps as a single run with a 10% burn-in. The SC model was run for a larger number of steps than the DTA model as its population and transmission parameters took longer to converge. BEAST 1.8.3 is unable to run the SC model, unlike BEAST 2.4. BEAST 2.4 can run GMRF and DTA models but does not have a BEAUti interface to easily set up these models. BEAST 1.8.3. does have an interface for these models so was used for the DTA model.

### Model comparison

The SC and DTA models were used to estimate the amount of time spent in each host population (population parameters) and the amount of transmissions between the host populations (transmission parameters). However, the models’ raw outputs were not directly comparable, as the SC model’s implementation explicitly records transmissions along branches, whilst the DTA approach integrates and marginalizes over these transmissions and therefore does not record them in its output. Therefore, the relative amount of time (i.e. proportion) spent in each host population and the relative number of inter-population transmissions made up of each transmission were compared. The performance of the two models were compared using four parameters:

1. The proportion of outbreak simulations that a model included the known parameter within their 95% highest posterior density (HPD) intervals.
2. The mean squared error between a known parameter and a model’s mean estimates.
3. The size of a model’s 95% HPD intervals.
4. The correlation coefficient between a known parameter and a model’s mean estimates.

### DT160 outbreak

The DTA and SC models were used to analyze a previously-described salmonellosis outbreak in New Zealand caused by DT160 (11). 109 DT160 isolates from animal (n=74) and human (n=35) host populations over 14 years were investigated using the 793 core SNPs they shared.

### Scripts

The in-house scripts used in this study are available from GitHub (https://github.com/samuelbloomfield/Scripts-for-outbreak-simulations).

## Results

### Model consistency

There was some variation in the DTA and SC models’ population and transmission mean estimates for the same simulated outbreak that was randomly sampled ten times (Fig 2). The SC model’s 95% HPD intervals included known population parameters more frequently, whilst the DTA model’s 95% HPD intervals included known transmission parameters more frequently.

**Fig 2.**
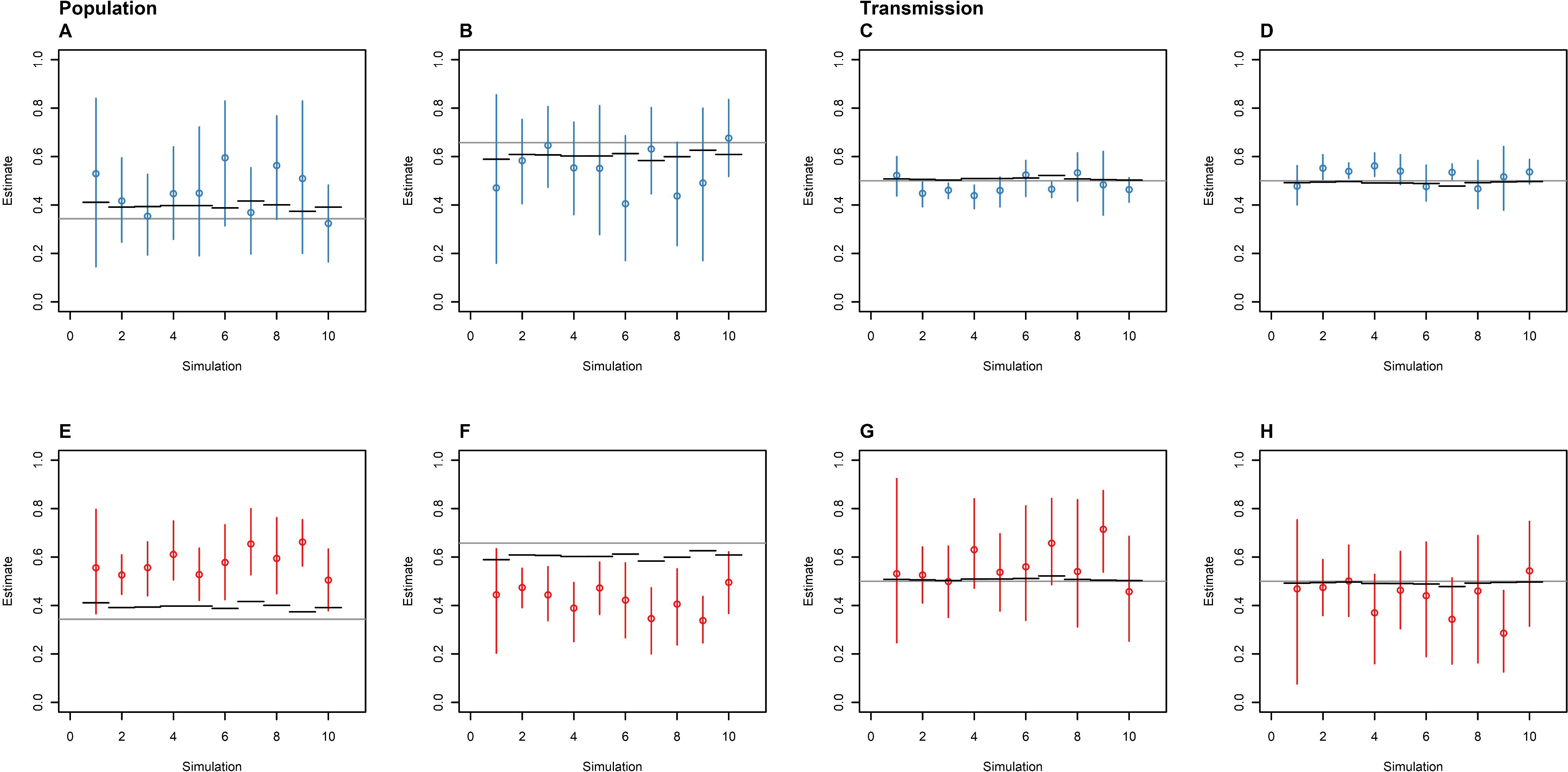
The proportion of time spent in the animal (A and E) and human (B and F) host populations, and the proportion of inter-population transmissions made up of animal-to-human (C and G) and human-to-animal (D and H) transmissions as estimated by the SC (blue: A-D) and DTA (red: E-F) models, for 10 random samples of the same simulated outbreak. The circles represent the mean, the error bars represent the 95% HPD interval, the black horizontal lines represent the known parameters for the sampled outbreaks, and the grey horizontal lines represent the known parameters for the entire outbreak.

The outbreak transmission tree was the same for the ten samples, as these samples were taken from the same simulated outbreak. However, the samples consisted of different animal and human isolates, such that when the outbreak transmission tree was simplified to only include nodes and branches common to these isolates, there was some variation in the time spent in animal and human populations, and the number of transmissions between these populations between samples. The known parameters were taken from the ten sampled transmission trees, not the entire outbreak transmission tree, resulting in slight differences in the known parameters between the ten samples. This is true for other analyses below that sampled the same outbreak multiple times. Some of the outbreaks investigated in this outbreak consisted of hundreds of thousands of infected animals and humans (S1 Appendix), leaving large outbreak transmission trees that required large time periods to calculate the number of transmissions and time spend in the populations. The small amount of variation in the sampled transmission trees and the outbreak transmission tree for this dataset suggests that the sampled transmission tree parameters are representative of the outbreak transmission tree parameters.

### Sample size

The DTA and SC models were affected by variation in sample size for the same simulated outbreak differently. Increased sample sizes were associated with smaller 95% HPD intervals and more accurate and extreme mean population estimates by the SC model up to 100 samples. After this point, increased sample sizes had little effect on the precision, extremity or accuracy of the model’s mean population estimates (Fig 3). The DTA model’s mean population estimates were more precise than the SC model’s. Sample size had no effect on their accuracy but decreased the size of their 95% HPD intervals. The accuracy of the SC and DTA models’ mean transmission estimates and their 95% HPD intervals displayed some variation, but there were no trends with sample size.

**Fig 3.**
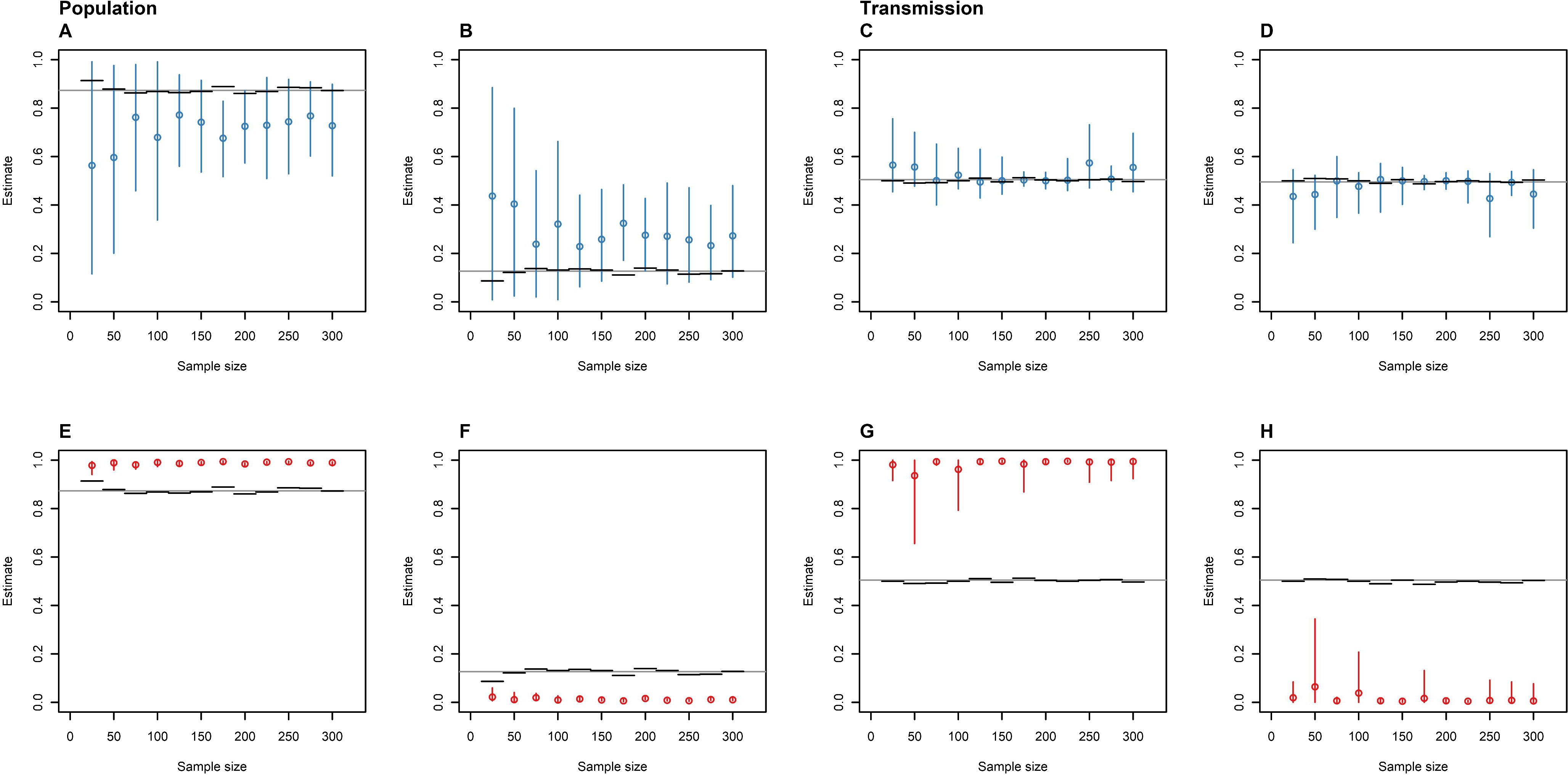
The proportion of time spent in the animal (A and E) and human (B and F) host populations, and the proportion of inter-population transmissions made up of animal-to-human (C and G) and human-to-animal (D and H) transmissions as estimated by the SC (blue: A-D) and DTA (red: E-F) models versus the number of isolates sampled from the same outbreak. The circles represent the mean, the error bars represent the 95% HPD interval, the black horizontal lines represent the known parameters for the sampled outbreaks, and the grey horizontal lines represent the known parameters for the entire outbreak.

### Disproportionate sampling

The DTA and SC models responded to variation in sample proportions for the same simulated outbreak differently. The DTA model’s mean estimates showed a much stronger positive correlation with the proportion of isolates sampled from each host population than the SC models’ mean estimates (Fig 4). The DTA model’s mean estimates displayed a sigmoid-like association with the proportion of isolates sampled from each host population (Fig 5).

**Fig 4.**
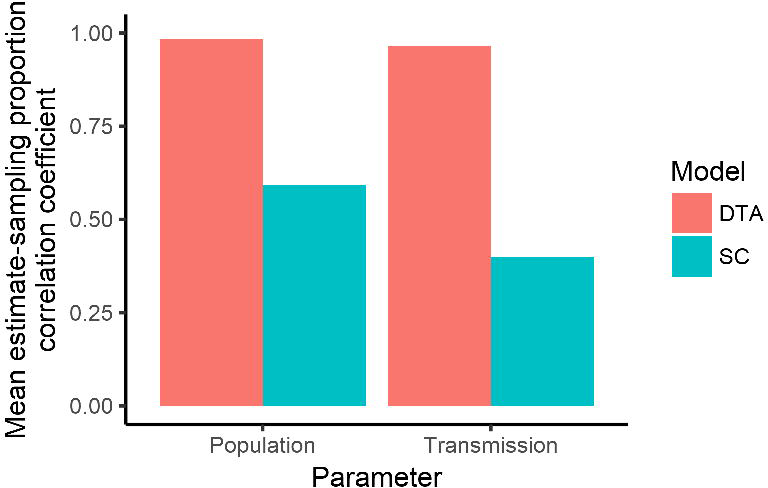
Bar graph of the correlation coefficients between the models’ mean estimates and the proportion of sampled isolates that are animal or human hosts for the same outbreak that was disproportionately sampled.

**Fig 5.**
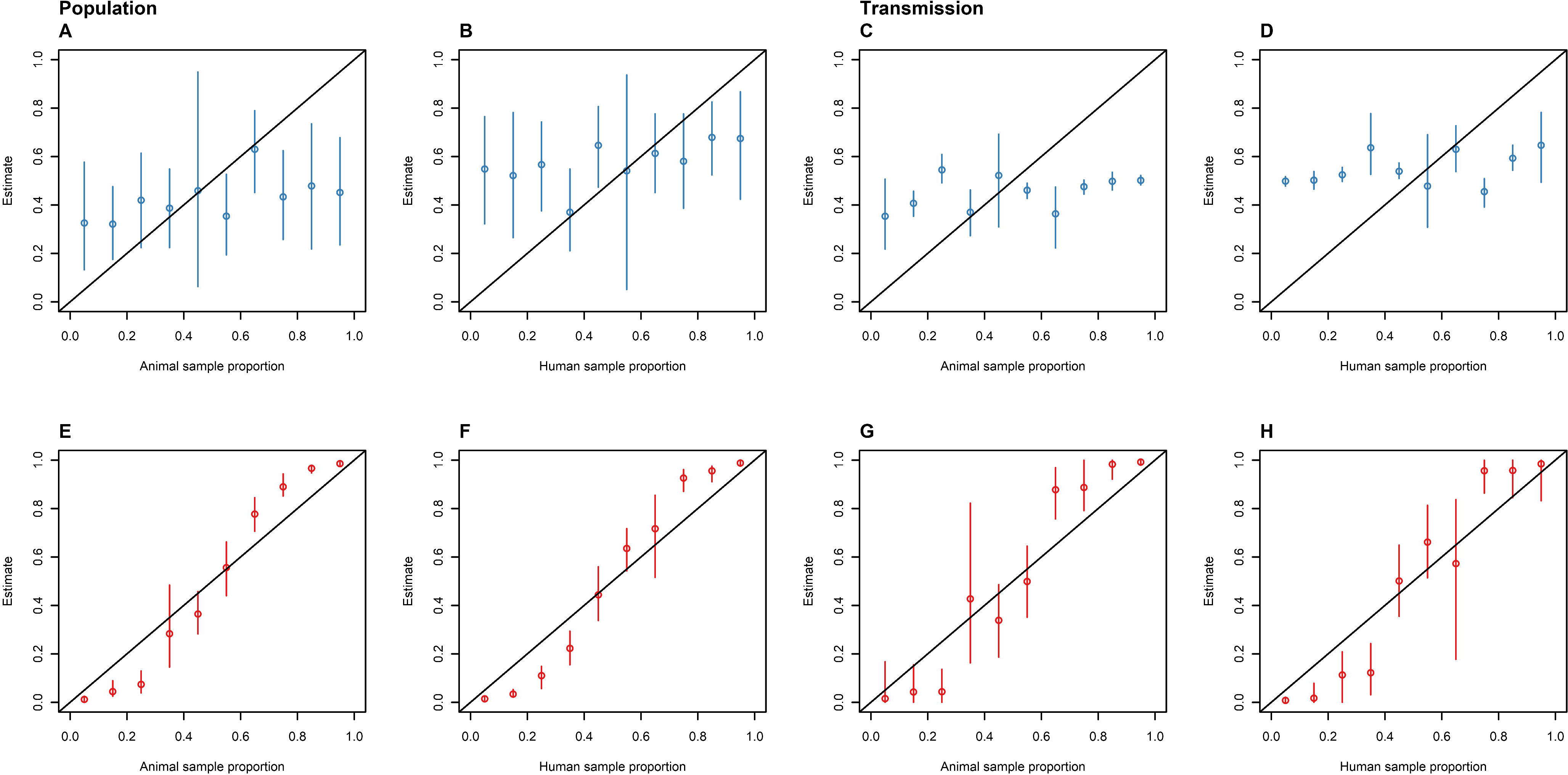
The proportion of time spent in the animal (A and E) and human (B and F) host populations, and the proportion of inter-population transmissions made up of animal-to-human (C and G) and human-to-animal (D and H) transmissions as estimated by the SC (blue: A-D) and DTA (red: E-F) models versus the proportion of sampled isolates that are animal (A, C, E and G) and human (B, D, F and H) for the same outbreak that was disproportionately sampled. The diagonal line represents accurate parameter estimates of the sampled outbreaks, the dots represent the mean, and the error bars represent the 95% HPD interval.

### Multiple variable simulations

The DTA and SC models showed different associations between known and estimated parameters when 100 isolates were randomly sampled from each of the 23 simulated outbreaks. The SC model predicted a larger proportion of known population and transmission parameters within its 95% HPD interval compared to the DTA model (Fig 6). However, its mean 95% HPD interval sizes were larger and the DTA model’s mean estimates showed a stronger positive correlation with the known parameter values than the SC model’s mean estimates. Both models had similar mean squared errors between the known parameters and the models’ mean estimates. However, the SC model’s mean population estimates were all within the 0.2-0.8 interval and its mean transmission rates were all within the 0.35-0.65 interval, whilst the DTA models had mean estimates that lay outside of these ranges (Fig 7).

**Fig 6.**
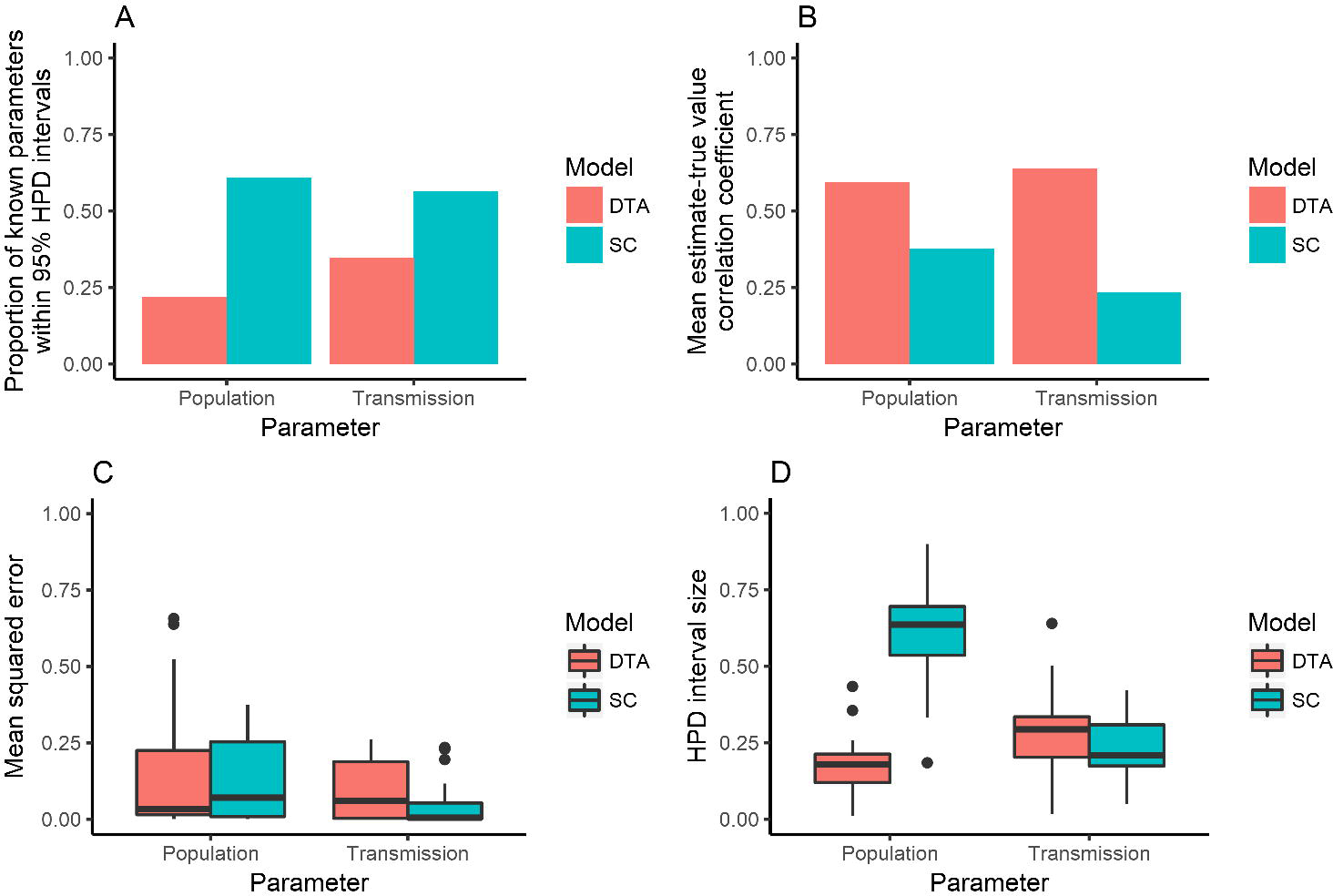
The proportion of outbreak simulations that the models included the known parameter within their 95% highest posterior density (HPD) intervals (A); the correlation coefficient between known parameters and the models’ mean estimates (B); the mean squared error between known parameters and the models’ mean estimates (C); and the size of the models’ 95% HPD intervals (D), for the population and transmission estimates made by the DTA (red) and SC (blue) models for 23 randomly-sampled simulated outbreaks that 100 isolates were randomly sampled from.

**Fig 7.**
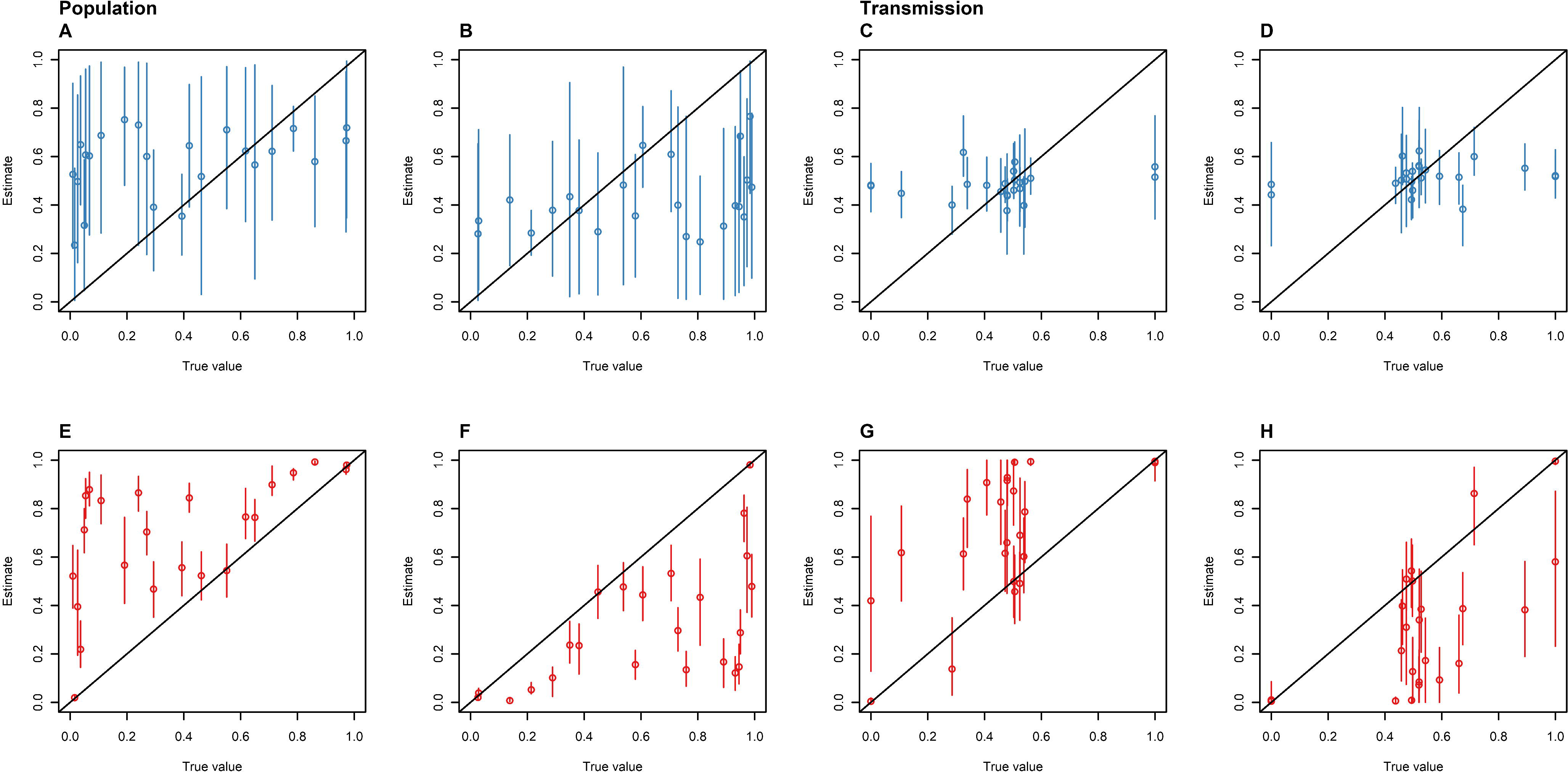
The proportion of time spent in the animal (A and E) and human (B and F) host populations, and the proportion of inter-population transmissions made up of animal-to-human (C and G) and human-to-animal (D and H) transmissions as estimated by the SC (blue: A-D) and DTA (red: E-F) models versus the true parameters for 23 simulated outbreaks that 100 isolates were randomly sampled from. The diagonal line represents accurate parameter estimates of the sampled outbreaks, the dots represent the mean, and the error bars represent the 95% HPD interval.

The phylogenetic trees produced by the DTA and SC models for the 23 simulated outbreaks poorly reflected the sampled transmission trees (Fig 8). The DTA model was unable to detect transmissions along branches in the transmission trees. The SC model could identify transmissions along branches, but often over-estimated the amount of transmissions compared to the true transmission tree. In the example given, the SC model predicted that ‘*Salmonella*’ was predominantly in the animal (red) population, as indicated by the predominantly red branches, but that coalescent events primarily occurred in the human (blue) population. This was common for most of the *maximum a priori* trees produced by the SC model, where the population that was estimated to have a smaller effective population size would be where the coalescent events took place, whilst the population with the estimated larger effective population size would predominate the branches. The phylogenetic trees in Fig 8 represent the most likely trees estimated using the DTA and SC models for one simulated outbreak, not the variation amongst each model, as each model estimated thousands of phylogenetic trees.

**Fig 8.**
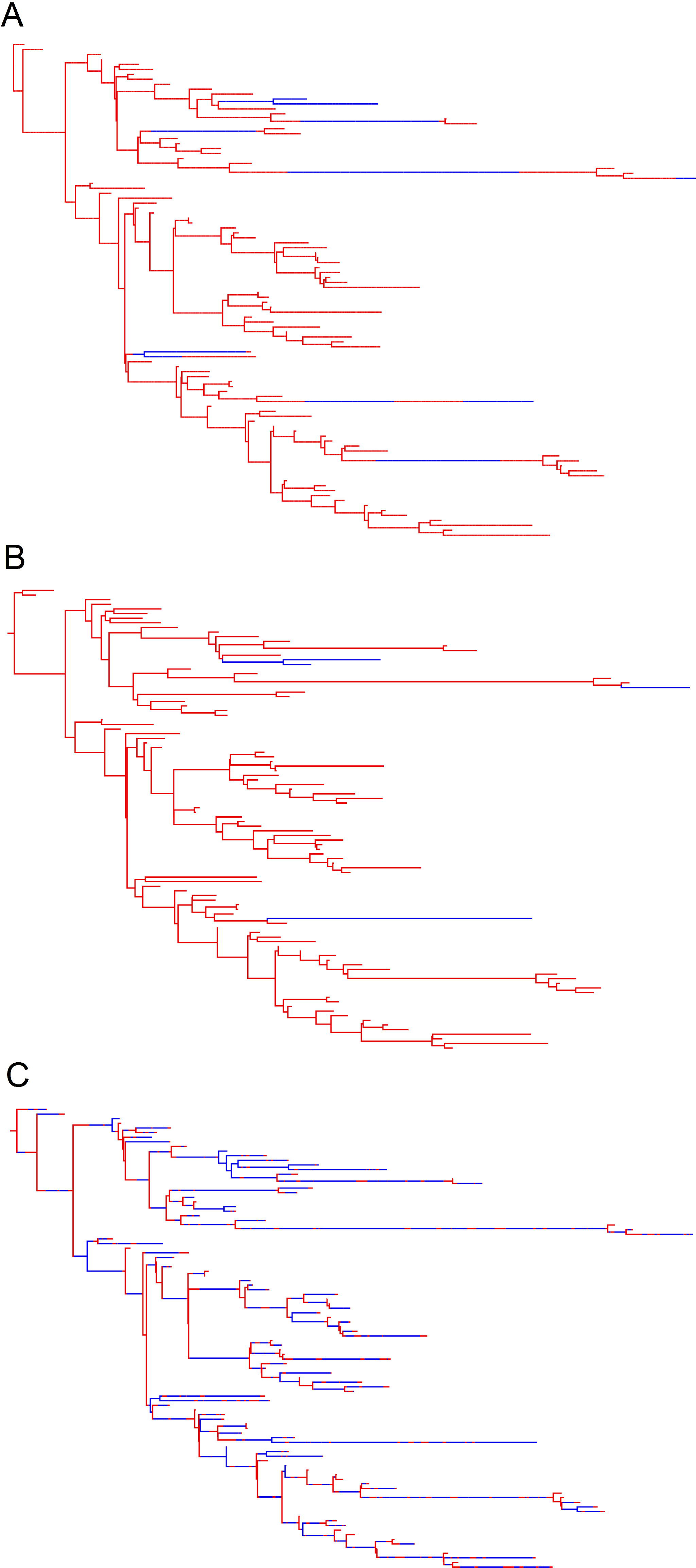
Sampled transmission tree (A), maximum clade credibility tree produced by the DTA model (B) and *maximum a posteriori* tree produced by the SC model (C), for one of the 23 simulated outbreaks that 100 isolates were randomly sampled from. The blue areas represent time spent in the human population and the red areas represent time spent in the animal population.

### Equal-time sampling

The DTA and SC models gave similar population and transmission estimates for the 23 simulated outbreaks with random (Fig 6–7) and equal-time sampling (Fig 9–10) of 100 isolates. Random sampling estimated more known parameters within its 95% HPD interval, but equal-time sampling had smaller mean squared errors between known parameters and the mean estimates, and smaller 95% HPD intervals. The SC and DTA models also estimated similar phylogenetic trees for simulated outbreaks that were sampled using random and equal-time sampling (Fig 11). This suggests that neither sampling method was more suitable for these ancestral state reconstruction models.

**Fig 9.**
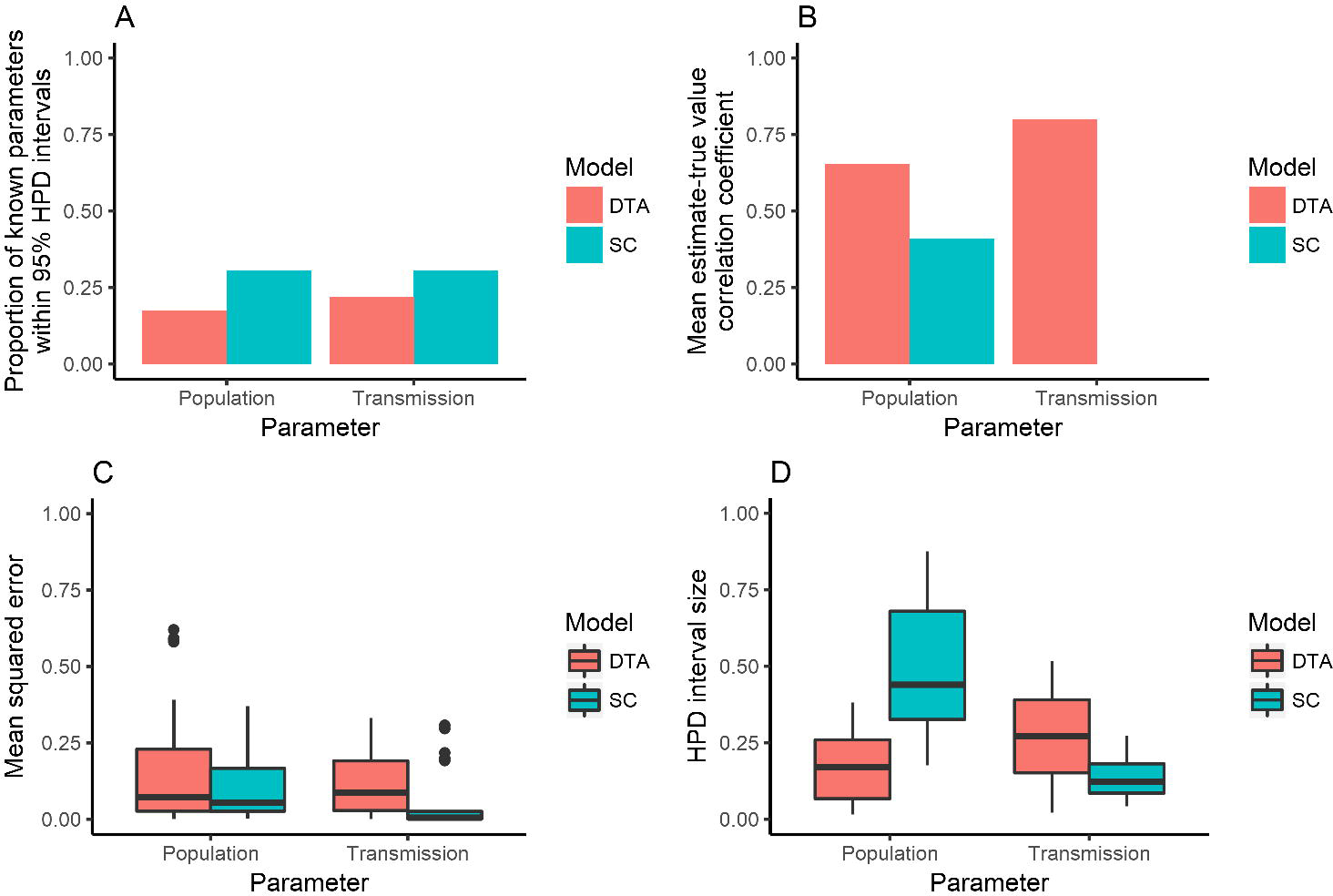
The proportion of outbreak simulations that the models included the known parameter within their 95% highest posterior density (HPD) intervals (A); the correlation coefficient between known parameters and the models’ mean estimates (B); the mean squared errors between known parameters and the models’ mean estimates (C), and the size of the models’ 95% HPD intervals (D), for the population and transmission estimates made by the DTA (red) and SC (blue) models for 23 simulated outbreaks that 100 isolates were sampled equally over time from.

**Fig 10.**
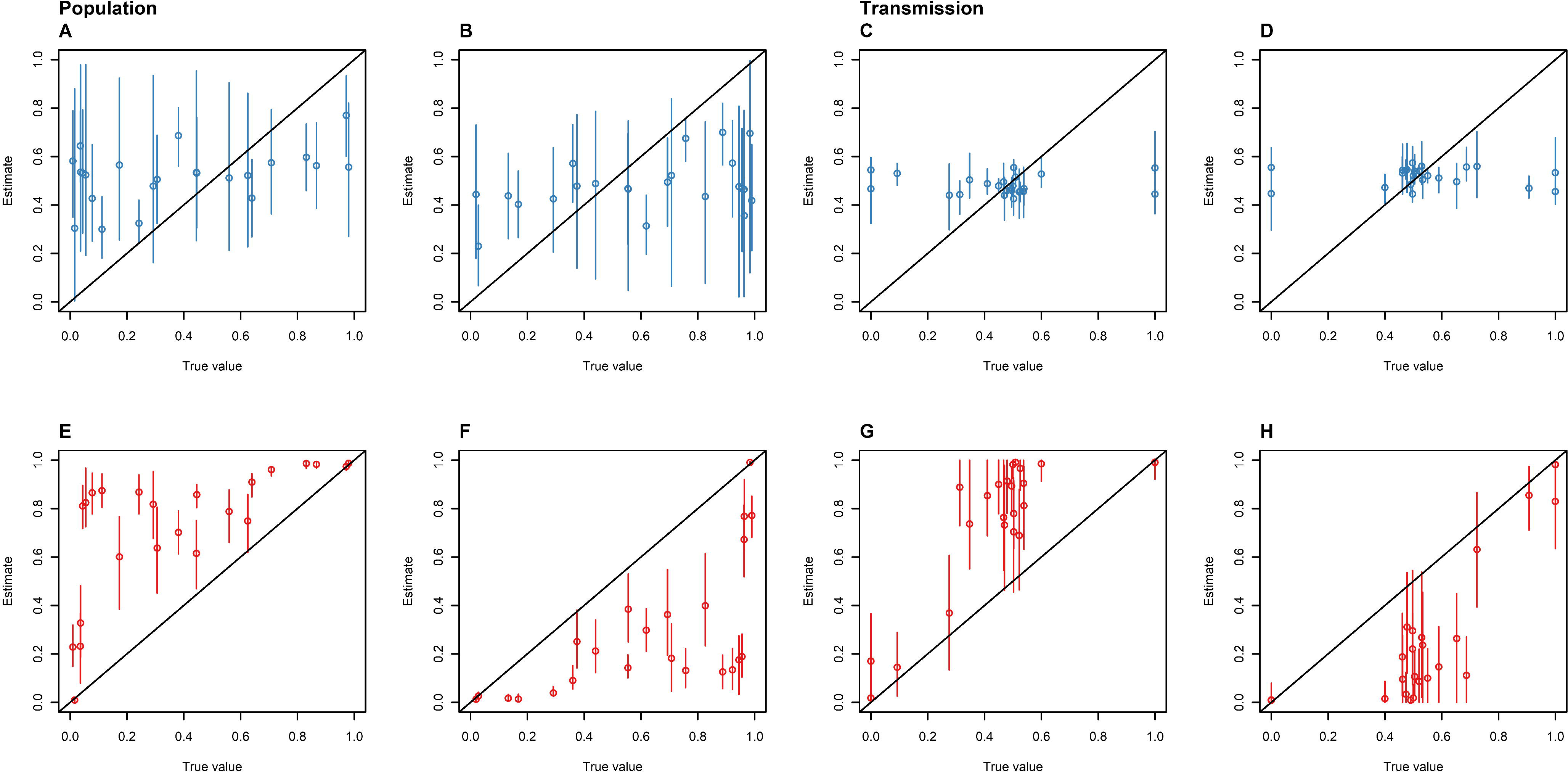
The proportion of time spent in the animal (A and E) and human (B and F) host populations, and the proportion of inter-population transmissions made up of animal-to-human (C and G) and human-to-animal (D and H) transmissions as estimated by the SC (blue: A-D) and DTA (red: E-F) models versus the true parameters for 23 simulated outbreaks that 100 isolates were sampled equally over time from. The diagonal line represents accurate estimates of the sampled outbreaks, the dots represent the mean, and the error bars represent the 95% HPD interval.

**Fig 11.**
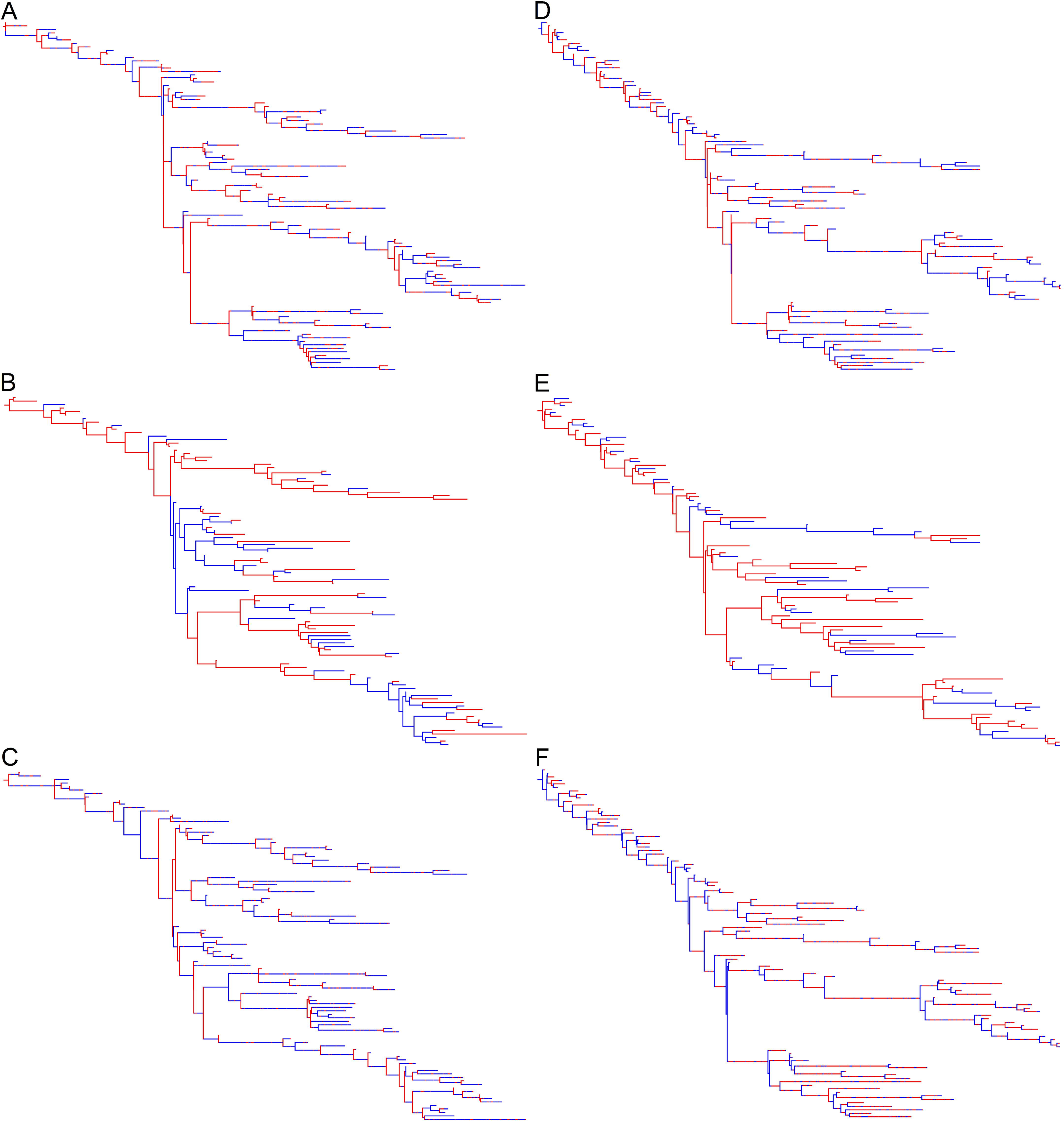
Sampled transmission tree (A and D), maximum clade credibility tree produced by the DTA model (B and E) and *maximum a posteriori* tree produced by the SC model (C and F), for one of the 23 simulated outbreaks that 100 isolates were sampled randomly (A-C) and equally over time (D-F). The blue areas represent time spent in the human population and the red areas represent time spent in the animal population.

### Equal intra-population transmission rates and infectious periods

The DTA and SC models provided more accurate estimates of population parameters for the 12 simulated outbreaks with equal intra-population transmission rates and infectious periods (EPTI) (Fig 12 and 13) than the 23 simulations where these parameters varied (Fig 6 and 7), with smaller mean squared errors, a higher proportion of known parameter within their 95% HPD intervals, and mean estimates that were more positively correlated with the known parameters. The DTA model’s mean population estimates displayed a sigmoid shape, similar to the simulated outbreak that was disproportionately sampled (Fig 5). On the other hand, the DTA and SC models gave less accurate transmission estimates for the 12 outbreaks with equal intra-population transmission rates and infectious periods between host populations than for the 23 simulations where these parameters varied, with larger mean squared errors, a lower proportion of known parameter within their 95% HPD intervals, and mean estimates that were less positively correlated or negative correlated with the known parameters.

**Fig 12.**
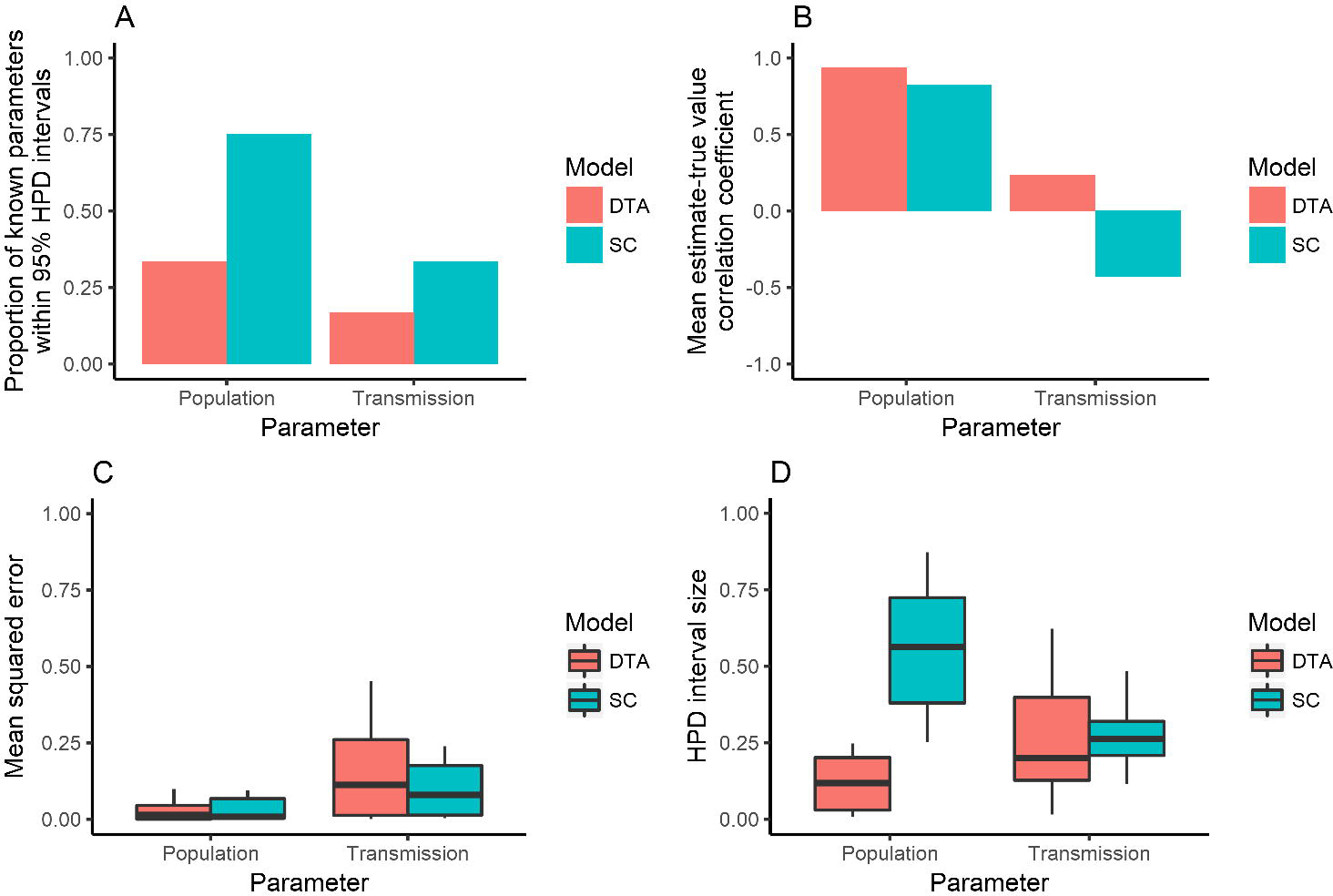
The proportion of outbreak simulations that the models included the known parameter within their 95% highest posterior density (HPD) intervals (A); the correlation coefficients between known parameters and the models’ mean estimates (B); the mean squared error between known parameters and the models’ mean estimates (C); and the size of the models’ 95% HPD intervals (D), for the population and transmission estimates made by the DTA (red) and SC (blue) models for 12 EPTI simulated outbreaks that 100 isolates were randomly sampled from.

**Fig 13.**
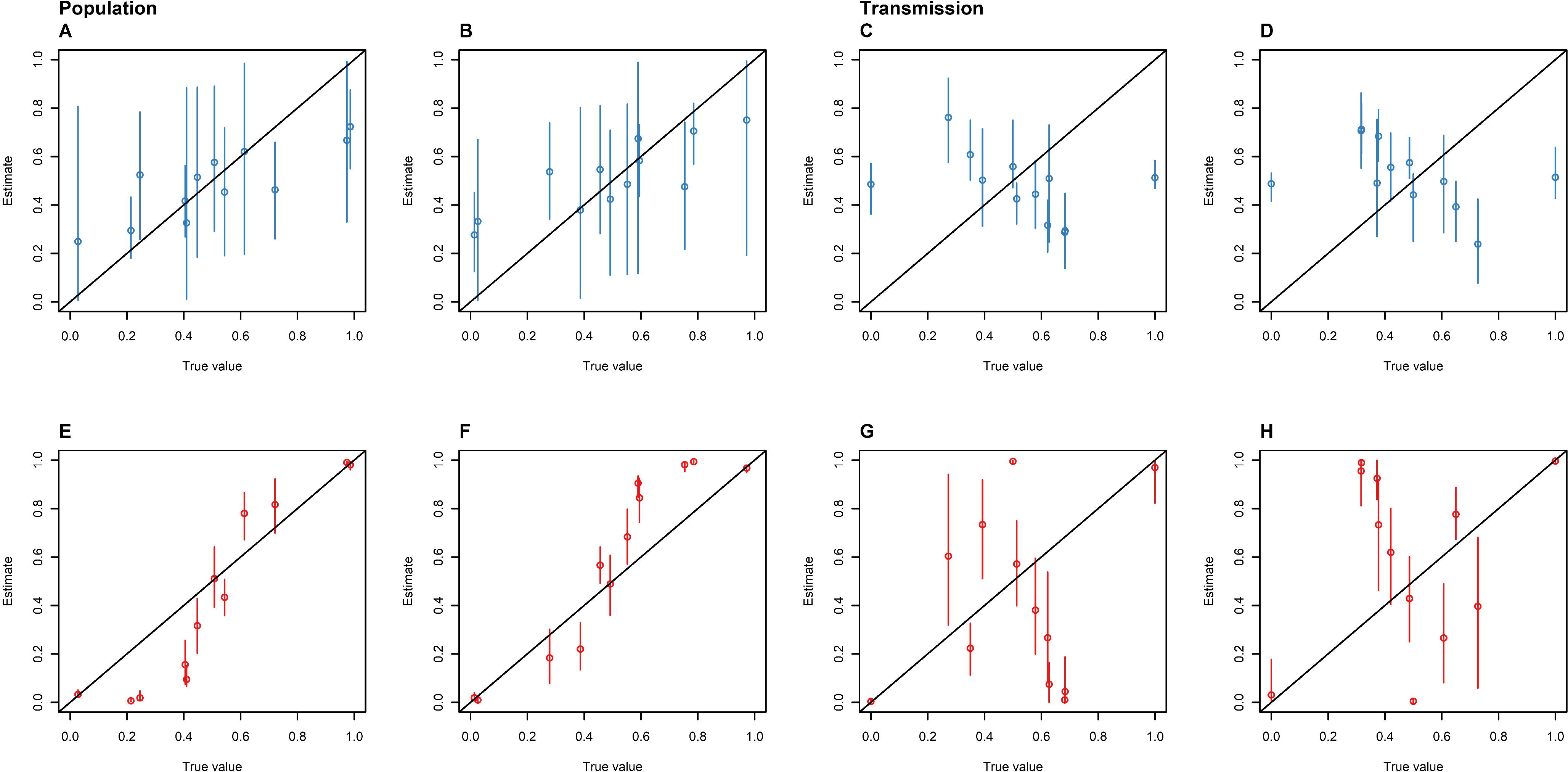
The proportion of time spent in the animal (A and E) and human (B and F) host populations, and the proportion of inter-population transmissions made up of animal-to-human (C and G) and human-to-animal (D and H) transmissions as estimated by the SC (blue: A-D) and DTA (red: E-F) models versus the true parameters for 12 EPTI simulated outbreaks that 100 isolates were randomly sampled from. The diagonal line represents accurate estimates of the sampled outbreaks, the dots represent the mean, and the error bars represent the 95% HPD interval.

The phylogenetic trees estimated for the 12 EPTI outbreaks (Fig 14) were like those of previous simulated outbreaks (Fig 8). They also demonstrated that the DTA model was unable to estimate ancestral branch states that were a different host population to daughter branches and tips. The SC model could estimate the state of ancestral branches that differed to the tips, but often estimated these branches inaccurately.

**Fig 14.**
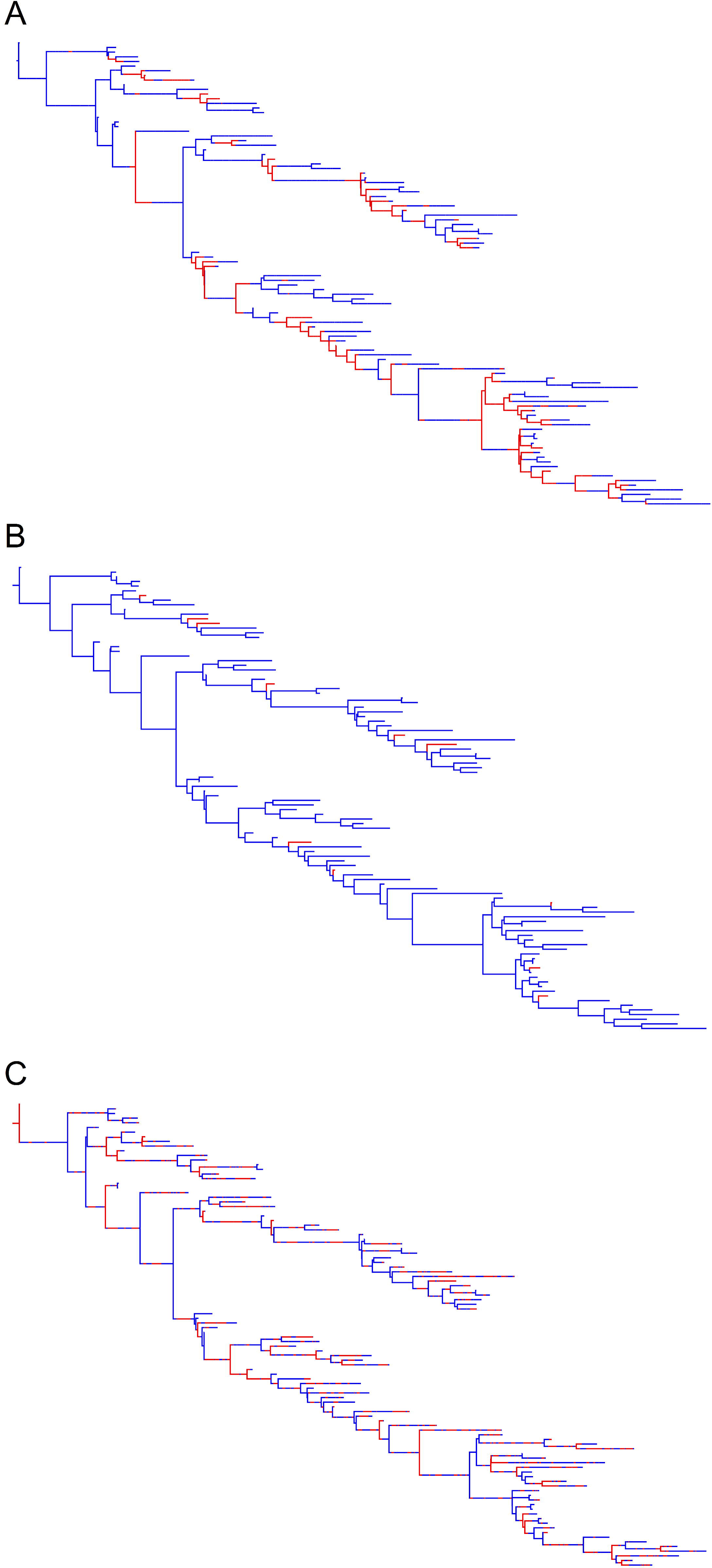
Sampled transmission tree (A), maximum clade credibility tree produced by the DTA model (B) and maximum a posteriori tree produced by the SC model (C), for a EPTI simulated outbreak that 100 isolates were randomly sampled from. The blue areas represent time spent in the human population and the red areas represent time spent in the animal population.

### Host sampling effect on the models’ estimates

To determine the effect of host sampling on the SC and DTA models’ estimates, the correlation coefficient between the proportion of samples isolated from each host population and the mean estimates for the simulated outbreaks were calculated (Fig 15; S1-S3 Fig). The DTA model’s mean population and transmission estimates were more positively correlated with the proportion of samples isolated from each population, than the SC model’s. The DTA model’s mean estimates displayed similar correlation coefficients for the 12 EPTI simulations and the 23 simulated outbreaks that were sampled randomly and equally over time, whilst the SC model’s estimates gave different correlation coefficients for these datasets.

**Fig 15.**
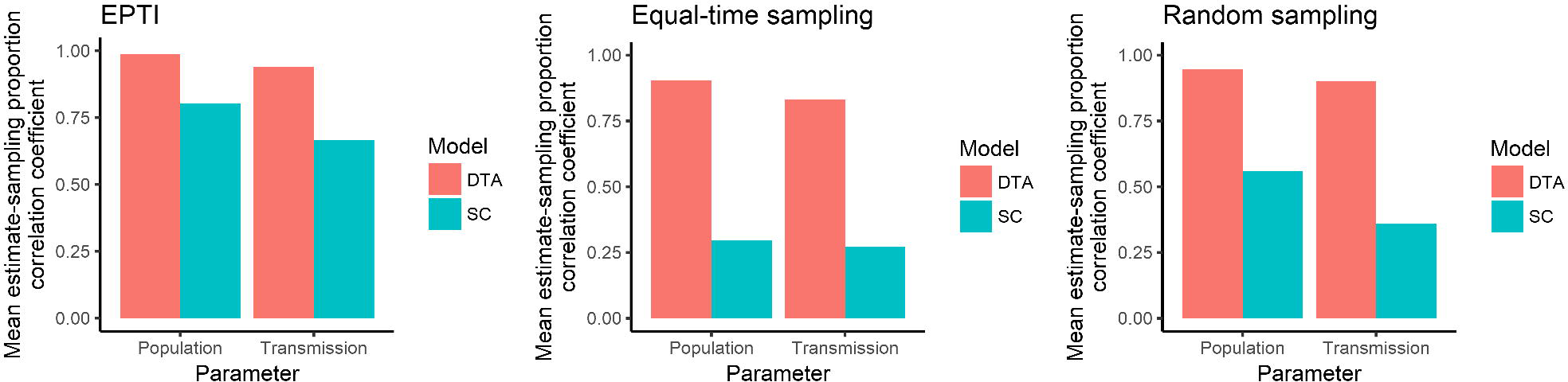
Bar graph of the correlation coefficients between the SC and DTA models’ mean estimates and the proportion of isolates sampled from each host population for 12 EPTI simulated outbreaks that 100 isolates were randomly sampled from, and 23 simulated outbreaks that 100 isolates were sampled randomly and equally over time.

To determine if the difference in sampling fraction could account for the DTA model’s estimates for the simulated outbreaks, the correlation coefficient between the proportion of samples isolated from each host and the known parameters were calculated (Fig 16; S4-S6 Fig). The known population parameters for the 12 EPTI simulated outbreaks and the sampling proportions were highly correlated, accounting for the more accurate estimates of these known parameters by the DTA model (Fig 13) compared to the known transmission parameters and other outbreak datasets where there was less correlation (Fig 7, 10, 13).

**Fig 16.**
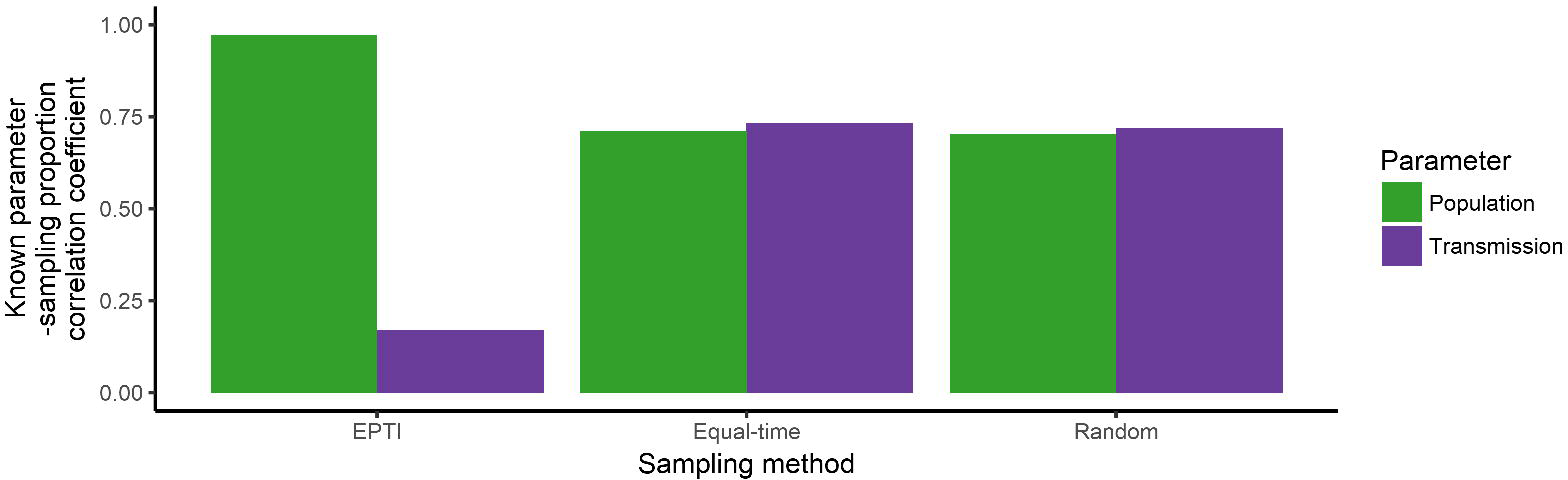
Bar graph of the correlation coefficients between the proportion of isolates sampled from each host population and the known population and transmission parameters for 12 EPTI simulated outbreaks that 100 isolates were randomly sampled from, and 23 simulated outbreaks that 100 isolates were sampled randomly and equally over time.

### DT160 outbreak

The SC and DTA models both predicted that DT160 spent most of the time in the animal host population over the course of the DT160 outbreak in New Zealand (Fig 17). However, the SC model predicted that there were relatively equal amounts of transmission between the animal and human host populations, whilst the DTA model predicted that there was a large amount of animal-to-human transmission and relatively less human-to-animal transmission. The phylogenetic trees estimated for the DT160 outbreak also displayed larger intervals between coalescent events later in the outbreak compared to the outbreaks simulated in this study (Fig 18).

**Fig 17.**
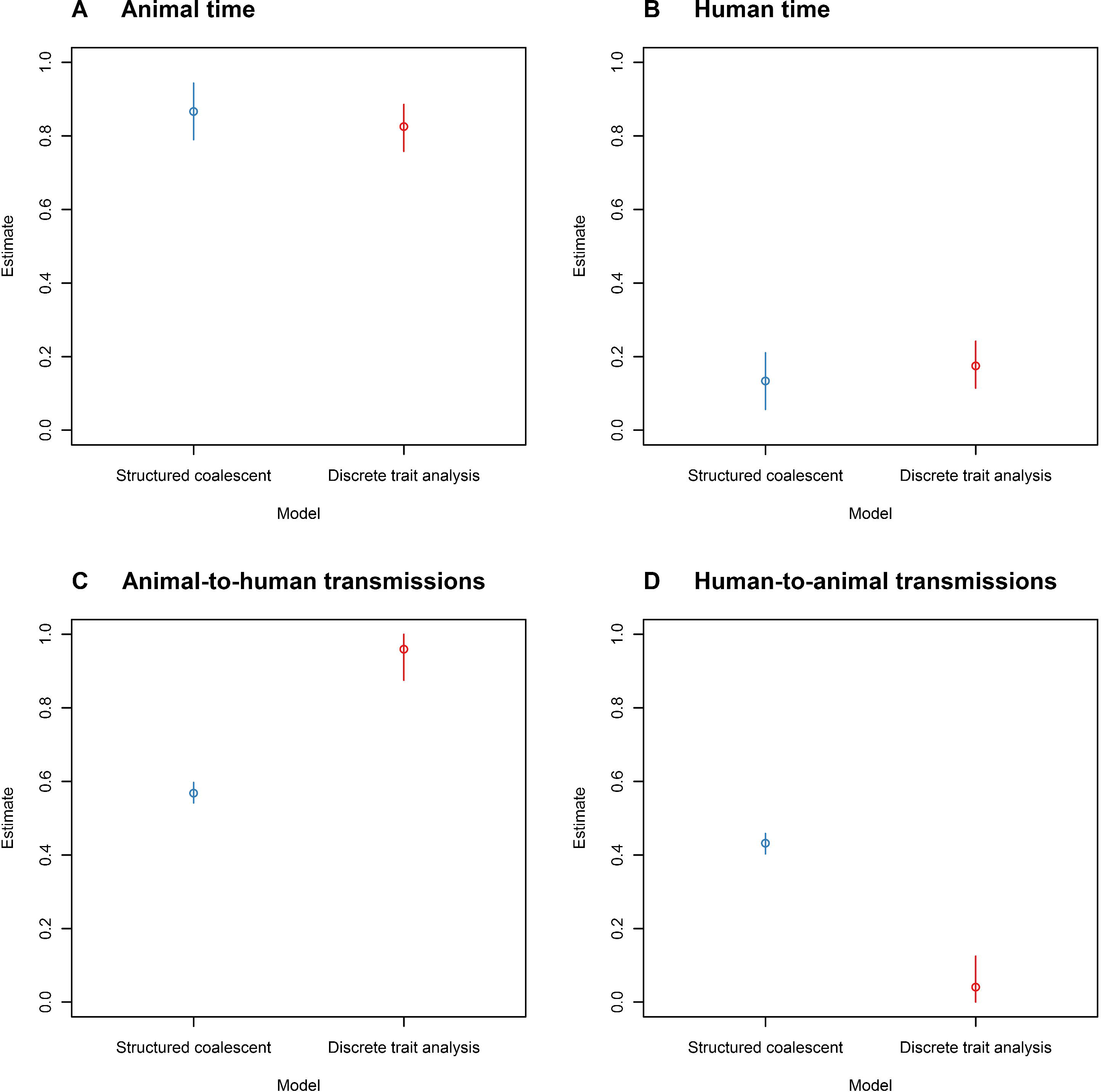
Estimates of the proportion of time spend in the animal (A) and human (B) host populations, and the proportion of inter-population transmissions made up of animal-to-human (C) and human-to-animal (D) transmissions for the DT160 outbreak, as estimated by the SC (blue) and DTA (red) models on 109 isolates. The circles represent the mean and the error bars represent the 95% HPD interval.

**Fig 18.**
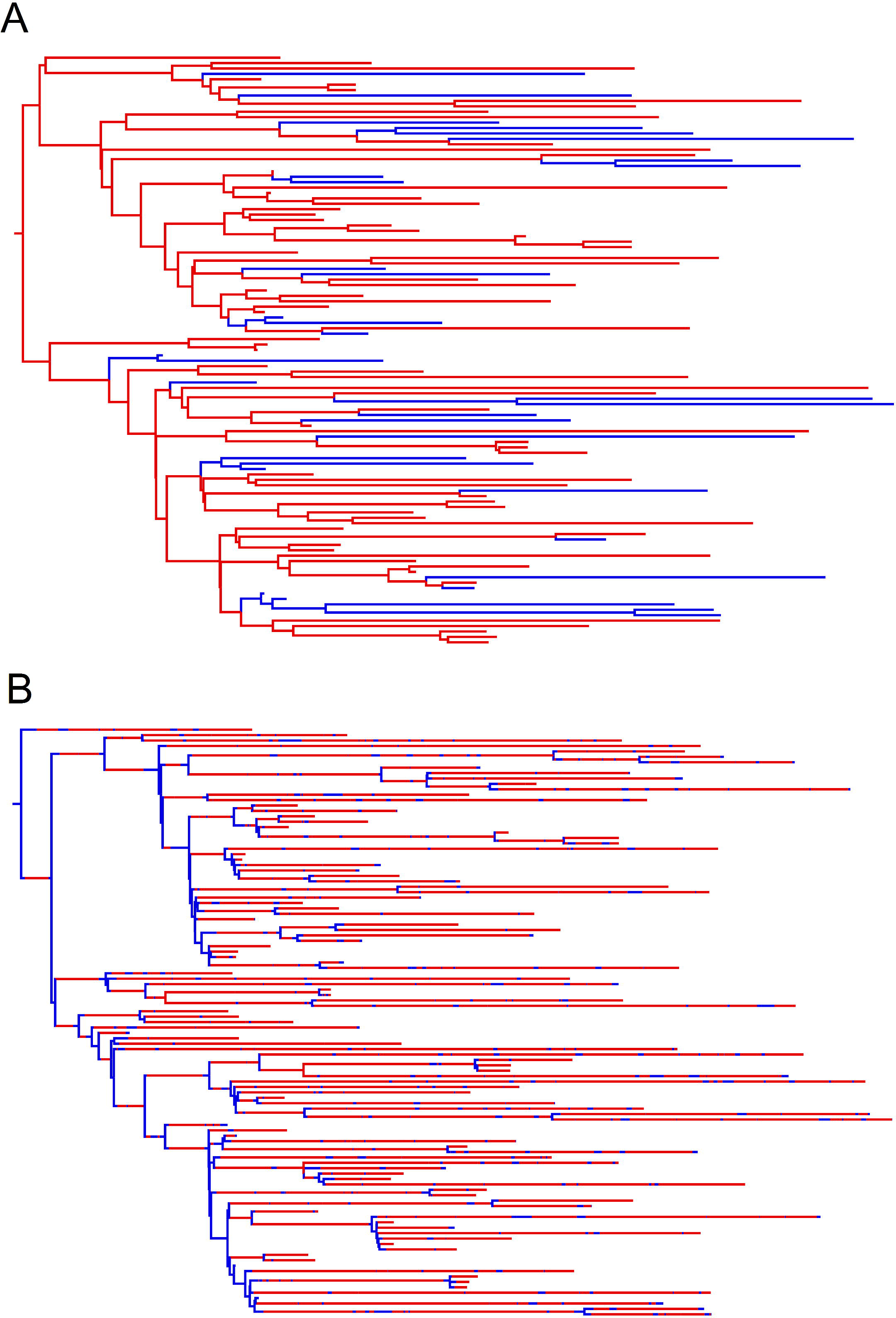
Maximum clade credibility tree produced by the DTA model (A) and *maximum a posteriori* tree produced by the SC model (B), based on 109 DT160 isolates.

## Discussion

The DTA and SC models are ancestral state reconstruction models that were designed to estimate the ancestral state of a group of organisms based on their evolutionary history (3,4). In this study we demonstrated using simulated outbreaks and a previously described salmonellosis outbreak that neither of these models could accurately estimate known population and transmission parameters for these outbreaks.

The DTA model assumes that the proportion of samples from each host population is proportional to its relative size (6). This is a problem for outbreaks involving multiple host populations, as the host populations may be sampled at different rates, resulting in samples disproportional to the number of individuals infected within each host population. The simulated outbreaks in this study were stratified by host population before random sampling in efforts to meet this assumption. However, differing intra-population transmission rates and infectious periods between the host populations resulted in inter-population transmission rates and length of times spent in host populations disproportionate to the number of individuals infected within each host population and thus the proportion of each population sampled.

This may explain why the DTA model consistently over-estimated the length of time in the animal host population and the number of animal-to-human transmissions for the initial 23 simulated outbreaks, as the human host populations of these outbreaks were simulated to have longer infectious periods than the animal host populations. This resulted in longer periods spent in the human host population and a larger number of human-to-animal transmissions relative to the number of humans sampled.

The DTA model appeared to estimate population parameters more accurately when the parameter was directly proportional to the number of isolates from each host population sampled. In these instances, the population estimates and simulated outbreak parameters shared a sigmoid-like relationship due to the model’s ancestral branch estimates: the DTA model usually predicts that all the ancestral branches are one host population, until the majority of the tips are another host population, where all the ancestral branches switch (11). The correct population parameters were also only estimated when simulating outbreaks with equal intra-population transmission rates and infectious periods, parameters that usually differ between *Salmonella* host populations (15,16). However, even in these instances the DTA model inaccurately estimated ancestral host population states and transmission parameters.

The SC model gave similar estimates for all the simulated outbreaks. It was poor at estimating simulated outbreaks known parameters, only accurately estimating them when they were within the range that it consistently estimated. The SC model’s inaccurate estimates are possibly due to the model’s assumption that the effective population size of the host populations were consistent throughout the outbreak (10), which does not apply to salmonellosis outbreaks whose effective population size varies over the course of the outbreak (11). There may be other reasons why the SC model was unable to detect a signal, but it is difficult to test for these without first accounting for the model’s effective population size assumption.

The inability of the SC and DTA models to accurately estimate salmonellosis outbreak parameters highlights the need for outbreak-specific models. These models would need to be able to take into consideration variable sampling between host populations, like the SC model, and changes in the effective population size, like the DTA model. In addition, they would need to be able to take into consideration variation in infectious periods and intra-population transmission rates.

The MASTER package of BEAST2 allowed many salmonellosis outbreaks to be simulated using the stochastic SIR model. The simulated outbreaks contained a large amount of variation in the amount of time spent in the animal and human host populations, but less variation in inter-population transmissions due to only simulating two host populations. Therefore, unequal transmission values were only simulated using one very high and one very low inter-population transmission value. This in part explains why the SC model was more likely to provide estimates that matched known simulation parameters because it always gave similar mean estimates around the 0.35-0.65 range, which most of the known transmission parameters for the simulated outbreaks were within. Further work with multiple host populations may help better understand these models’ application to salmonellosis outbreaks.

The DTA and SC models’ estimates of the DT160 outbreak underline some of the limitations of this study. The DTA model estimated that DT160 spent most of its time in the animal host population and that there was a larger amount of animal-to-human transmission than human-to-animal transmission, which is to be expected as the DTA model is affected by sample size and a larger number of animal isolates were analyzed than human isolates in the DT160 study. The SC model estimated similar amounts of animal-to-human transmission than human-to-animal transmission, which is also to be expected as our study shows it usually gives similar transmission rates between two host populations. However, the SC model estimated that DT160 spent over 90% of its time in the animal host population and less than 10% of its time in the human host population, outside the 20-80% range estimated for simulated outbreaks, and both models produced phylogenetic trees with larger distances between coalescent events towards the later part of the outbreak than simulated outbreaks. The effective population size affects the timing of coalescent events for randomly sampled individuals (17). This suggests that the DT160 outbreak had a much larger effective population size than any of the simulated outbreaks in this study. It also indicates that the SC model’s estimates maybe influenced by branch length. Simulations with larger effective population sizes are required to test this.

In conclusion, our comparison of applicability of the SC and DTA models to salmonellosis outbreaks between the known parameters of simulated outbreaks and the models’ estimates suggest neither model is appropriate for this analysis. Our findings highlight the need for outbreak-specific models that can also take into consideration intra-population transmission rates, infectious periods, disproportionate sampling and changes in the effective population size.

## Acknowledgements

We acknowledge the contribution of the New Zealand eScience Infrastructure (NeSI) high-performance computing facilities to the results of this research.

## S1 Appendix – Simulated outbreak parameters

**S1 Fig**. The proportion of time spent in the animal (A and E) and human (B and F) host populations, and the proportion of inter-population transmissions made up of animal-to-human (C and G) and human-to-animal (D and H) transmissions as estimated by the SC (blue: A-D) and DTA (red: E-F) models versus the proportion of samples made up of animal (A, C, E and G) and human (B, D, F and H) host populations for 12 EPTI simulated outbreaks that 100 isolates were randomly sampled from. The dots represent the mean, and the error bars represent the 95% HPD interval.

**S2 Fig**. The proportion of time spent in the animal (A and E) and human (B and F) host populations, and the proportion of inter-population transmissions made up of animal-to-human (C and G) and human-to-animal (D and H) transmissions as estimated by the SC (blue: A-D) and DTA (red: E-F) models versus the proportion of samples made up of animal (A, C, E and G) and human (B, D, F and H) host populations for 23 simulated outbreaks that 100 isolates were randomly sampled from. The dots represent the mean, and the error bars represent the 95% HPD interval.

**S3 Fig**. Scatterplots of the proportion of time spent in the animal (A and E) and human (B and F) host populations, and the proportion of inter-population transmissions made up of animal-to-human (C and G) and human-to-animal (D and H) transmissions as estimated by the SC (blue: A-D) and DTA (red: E-F) models versus the proportion of samples made up of animal (A, C, E and G) and human (B, D, F and H) host populations for 23 simulated outbreaks that 100 isolates were sampled equally over time from. The dots represent the mean, and the error bars represent the 95% HPD interval.

**S4 Fig**. The proportion of samples made up of animal (A and C) and human (B and D) host populations, versus the known population (A and B) and transmission (C and D) parameters for 12 EPTI simulated outbreaks that 100 isolates were randomly sampled from.

**S5 Fig**. The proportion of samples made up of animal (A and C) and human (B and D) host populations, versus the known population (A and B) and transmission (C and D) parameters for 23 simulated outbreaks that 100 isolates were randomly sampled from.

**S6 Fig**. The proportion of samples made up of animal (A and C) and human (B and D) host populations, versus the known population (A and B) and transmission (C and D) parameters for 23 simulated outbreaks that 100 isolates were sampled equally over time from.

