## Supplementary material for "Investigation of the validity of two Bayesian ancestral state reconstruction models for estimating *Salmonella* transmission during outbreaks": S1 Appendix

**S1 Appendix - Simulated outbreak parameters**

The outbreaks simulated in this study were designed to represent possible salmonellosis outbreaks in New Zealand. This was achieved by using host population and infectious period parameters of humans and animals in New Zealand.

**Population size**

The human population in New Zealand increased linearly from 3.8 million in 1998 to 4.6 million in 2015 (1). The susceptible human host population size was taken from this statistic (S1 Table).

Cows and chickens are the primary sources of salmonellosis outbreaks (2). The number of cows in New Zealand fluctuated from 7.7-9.3 million in New Zealand from 1971-1999, whilst the number of chickens in New Zealand fluctuated from 18.3-21.9 million from 2002-2014 in New Zealand (3). Population records outside these time periods are scant for these animals. The susceptible host animal population size was taken by summing up these populations (S1 Table).

**S1 Table.** Population sizes for 23 simulated outbreaks

|  | Population size | |  |  |
| --- | --- | --- | --- | --- |
| Outbreak | Initial susceptible animal | Initial susceptible human | Total infected animal | Total infected human |
| 1 | 2.60E+07 | 4.50E+06 | 1.84E+02 | 4.25E+03 |
| 2 | 2.50E+07 | 4.20E+06 | 3.70E+03 | 5.56E+03 |
| 3 | 2.60E+07 | 3.80E+06 | 4.51E+03 | 6.40E+02 |
| 4 | 3.00E+07 | 4.00E+06 | 5.41E+03 | 6.91E+03 |
| 5 | 3.20E+07 | 4.60E+06 | 5.47E+03 | 7.97E+03 |
| 6 | 2.50E+07 | 4.20E+06 | 6.74E+03 | 3.25E+03 |
| 7 | 2.95E+07 | 3.90E+06 | 7.81E+03 | 4.71E+03 |
| 8 | 2.50E+07 | 4.20E+06 | 8.98E+03 | 9.63E+03 |
| 9 | 2.50E+07 | 4.20E+06 | 9.58E+03 | 1.87E+04 |
| 10 | 2.50E+07 | 4.20E+06 | 1.04E+04 | 1.05E+04 |
| 11 | 2.90E+07 | 3.85E+06 | 1.22E+04 | 4.46E+03 |
| 12 | 2.50E+07 | 4.20E+06 | 1.30E+04 | 1.06E+04 |
| 13 | 2.50E+07 | 4.60E+06 | 1.34E+04 | 3.51E+02 |
| 14 | 2.70E+07 | 4.50E+06 | 1.49E+04 | 5.91E+03 |
| 15 | 2.50E+07 | 4.20E+06 | 1.62E+04 | 1.55E+04 |
| 16 | 2.50E+07 | 4.20E+06 | 1.90E+04 | 1.85E+04 |
| 17 | 2.50E+07 | 4.20E+06 | 1.91E+04 | 9.86E+03 |
| 18 | 2.50E+07 | 4.20E+06 | 2.28E+04 | 1.53E+04 |
| 19 | 2.50E+07 | 4.20E+06 | 2.57E+04 | 1.08E+04 |
| 20 | 2.70E+07 | 4.50E+06 | 2.57E+04 | 1.15E+04 |
| 21 | 2.50E+07 | 4.20E+06 | 3.03E+04 | 5.15E+03 |
| 22 | 3.10E+07 | 3.90E+06 | 1.01E+05 | 4.26E+03 |
| 23 | 2.50E+07 | 4.60E+06 | 3.16E+05 | 1.21E+04 |

**Transmission rates**

A large range of transmission rates between the animal and human populations were used as these were the parameters being estimated in this study (S2 Table).

**S2 Table.** Transmission and removal rates for 23 simulated outbreaks

|  | Removal rate  γ (year^-1^ I^-1^) | | Transmission rate  β (year^-1^ I^-1^ S^-1^) | | | |
| --- | --- | --- | --- | --- | --- | --- |
| Outbreak | Animal | Human | Animal-to-animal | Human-to-human | Animal-to-human | Human-to-animal |
| 1 | 15 | 10 | 8.00E-07 | 2.22E-06 | 1.00E-09 | 5.00E-09 |
| 2 | 35 | 13 | 1.99E-07 | 2.97E-06 | 4.95E-07 | 2.97E-07 |
| 3 | 16 | 13 | 6.20E-07 | 1.00E-09 | 6.20E-07 | 1.00E-10 |
| 4 | 58 | 11 | 1.35E-07 | 2.70E-06 | 1.35E-07 | 2.70E-07 |
| 5 | 14 | 10 | 1.00E-09 | 2.18E-06 | 1.00E-09 | 2.18E-07 |
| 6 | 28 | 12 | 7.80E-07 | 1.47E-06 | 1.58E-06 | 2.94E-07 |
| 7 | 22 | 10 | 5.65E-07 | 1.43E-07 | 1.25E-07 | 8.43E-07 |
| 8 | 35 | 13 | 6.50E-07 | 2.34E-06 | 2.34E-06 | 2.60E-07 |
| 9 | 35 | 13 | 6.72E-07 | 1.63E-06 | 2.04E-06 | 5.49E-07 |
| 10 | 35 | 13 | 7.71E-07 | 1.54E-06 | 4.32E-06 | 2.26E-07 |
| 11 | 52 | 10.5 | 1.85E-07 | 3.00E-07 | 6.43E-07 | 3.42E-07 |
| 12 | 28 | 12 | 8.24E-07 | 1.55E-06 | 2.47E-06 | 1.55E-07 |
| 13 | 14 | 13 | 5.60E-07 | 2.50E-06 | 5.00E-09 | 1.00E-09 |
| 14 | 54 | 11.5 | 1.02E-07 | 2.04E-06 | 1.02E-06 | 1.02E-06 |
| 15 | 35 | 13 | 1.05E-06 | 9.50E-07 | 5.70E-06 | 1.33E-07 |
| 16 | 35 | 13 | 1.10E-06 | 8.23E-07 | 6.13E-06 | 1.15E-07 |
| 17 | 28 | 12 | 6.12E-07 | 1.84E-06 | 1.23E-06 | 4.29E-07 |
| 18 | 28 | 12 | 9.88E-07 | 9.88E-07 | 2.96E-06 | 9.88E-08 |
| 19 | 28 | 12 | 9.38E-07 | 9.38E-07 | 1.88E-06 | 1.88E-07 |
| 20 | 43 | 12.5 | 1.86E-07 | 1.86E-06 | 1.34E-06 | 9.30E-07 |
| 21 | 35 | 13 | 1.22E-06 | 5.00E-07 | 1.17E-06 | 4.00E-07 |
| 22 | 43 | 12.5 | 1.38E-06 | 2.07E-06 | 1.66E-07 | 1.38E-07 |
| 23 | 58 | 12 | 2.33E-06 | 2.50E-06 | 2.00E-08 | 1.00E-08 |

**Infectious periods**

The infectious period values used for the animal and human host populations were estimated from studies that measured the average length of *Salmonella* excretion after exposure. Buchwald and Blaser (4) reviewed 32 articles on human non-typhoid *Salmonella* excretion and found that patients excreted *Salmonella* for approximately 5 weeks. Murase et al. (5) investigated an outbreak of *Salmonella* *enterica* serovar Typhimurium in Japan and found that on average asymptomatic and symptomatic patients shed *Salmonella* for 4 weeks. The papers that Buchwald and Blaser reviewed focused on symptomatic salmonellosis patients, whilst Murase et al. considered asymptomatic patients that excreted *Salmonella* for a shorter length of time. This may explain why Murase et al. predicted a lower average *Salmonella* excretion rate than Buchwald and Blaser. However, Murase et al.'s study only represents a single outbreak. Therefore, the human *Salmonella* infectious period was taken from both studies (S3 Table).

The animal infectious period was estimated using studies on cow and chicken excretion periods. Gast et al. (6) exposed hens to different *S. enterica* serovar Enteritidis strains under different living conditions and found that on average the hens excreted *S.* Enteritidis for 2.0 weeks in conventional cages and 1.5 weeks in enriched colony cages. Barrow et al. (7) exposed multiple chicken lines to a strain of *S.* Typhimurium and a strain of *S.* Enteritidis and found that the average excretion varied between the strains and chicken lines (0.9-3.8 weeks). Alexander et al. (8) investigated two *S.* Typhimurium outbreaks in cows and found they shed *Salmonella* for 1.3-2 weeks. A broad range of animal infectious period values (0.9-3.8 weeks) were included to account for the large amount of variation in these studies (S2 Table).

**Equal intra-transmission rates and infectious periods**

Twelve outbreaks were simulated with identical infectious periods and intra-transmission rates between the host populations (EPTI), but inter-transmission rates and host population sizes that varied. For these outbreaks, all the transmission rates and host population sizes were within the ranges for the other simulated outbreaks (S1 and S2 Tables). However, the animal and human host populations shared infectious periods between 3.7-5.2 weeks (10-14 year^-1^ I^-1^) (S3 and S4 Tables). The intra-population transmission rates below appear to differ between the populations but are equal when the initial susceptible population size is taken into account, i.e. transmission rate is converted from β (year^-1^ I^-1^ S^-1^) to β (year^-1^ I^-1^).

**S3 Table.** Population sizes for 12 simulated EPTI outbreaks

|  | Population size | |  |  |
| --- | --- | --- | --- | --- |
| Outbreak | Initial susceptible animal | Initial susceptible human | Total infected animal | Total infected human |
| 24 | 2.60E+07 | 4.50E+06 | 2.69E+02 | 6.72E+03 |
| 25 | 2.70E+07 | 3.90E+06 | 1.20E+03 | 9.43E+03 |
| 26 | 3.10E+07 | 4.10E+06 | 1.26E+03 | 2.05E+04 |
| 27 | 2.55E+07 | 4.40E+06 | 1.38E+03 | 4.37E+03 |
| 28 | 2.75E+07 | 4.20E+06 | 2.34E+03 | 6.48E+03 |
| 29 | 2.80E+07 | 4.30E+06 | 3.52E+03 | 5.97E+03 |
| 30 | 2.50E+07 | 3.80E+06 | 4.66E+03 | 5.68E+03 |
| 31 | 3.00E+07 | 4.00E+06 | 4.94E+03 | 3.88E+03 |
| 32 | 2.90E+07 | 4.65E+06 | 6.38E+03 | 2.82E+03 |
| 33 | 2.60E+07 | 4.50E+06 | 9.22E+03 | 2.35E+03 |
| 34 | 3.20E+07 | 3.85E+06 | 9.52E+03 | 2.48E+02 |
| 35 | 2.50E+07 | 4.60E+06 | 1.02E+04 | 1.74E+02 |

**S4 Table.** Transmission and removal rates for 12 simulated EPTI outbreaks

|  | Removal rate  γ (year^-1^ I^-1^) | | Transmission rate  β (year^-1^ I^-1^ S^-1^) | | | |
| --- | --- | --- | --- | --- | --- | --- |
| Outbreak | Animal | Human | Animal-to-animal | Human-to-human | Animal-to-human | Human-to-animal |
| 24 | 10 | 10 | 3.85E-07 | 2.22E-06 | 1.00E-09 | 1.50E-09 |
| 25 | 12.5 | 12.5 | 4.04E-07 | 2.80E-06 | 3.00E-06 | 1.00E-08 |
| 26 | 13 | 13 | 3.84E-07 | 2.90E-06 | 6.00E-06 | 5.00E-09 |
| 27 | 10.5 | 10.5 | 3.36E-07 | 1.95E-06 | 1.50E-06 | 2.50E-08 |
| 28 | 11.5 | 11.5 | 3.36E-07 | 2.20E-06 | 1.50E-06 | 2.50E-08 |
| 29 | 13.5 | 13.5 | 3.95E-07 | 2.58E-06 | 1.00E-06 | 5.00E-08 |
| 30 | 12 | 12 | 3.57E-07 | 2.35E-06 | 1.00E-06 | 1.00E-07 |
| 31 | 11.5 | 11.5 | 3.60E-07 | 2.70E-06 | 7.00E-08 | 3.00E-08 |
| 32 | 10 | 10 | 3.17E-07 | 2.00E-06 | 5.00E-08 | 5.00E-08 |
| 33 | 14 | 14 | 5.11E-07 | 2.95E-06 | 2.50E-08 | 7.50E-08 |
| 34 | 11 | 11 | 3.43E-07 | 2.85E-06 | 1.00E-08 | 1.00E-07 |
| 35 | 14 | 14 | 5.60E-07 | 3.05E-06 | 4.00E-09 | 1.00E-09 |
