## Supplementary figures and images for "Investigation of the validity of two Bayesian ancestral state reconstruction models for estimating *Salmonella* transmission during outbreaks"

### Supplementary Figure 1

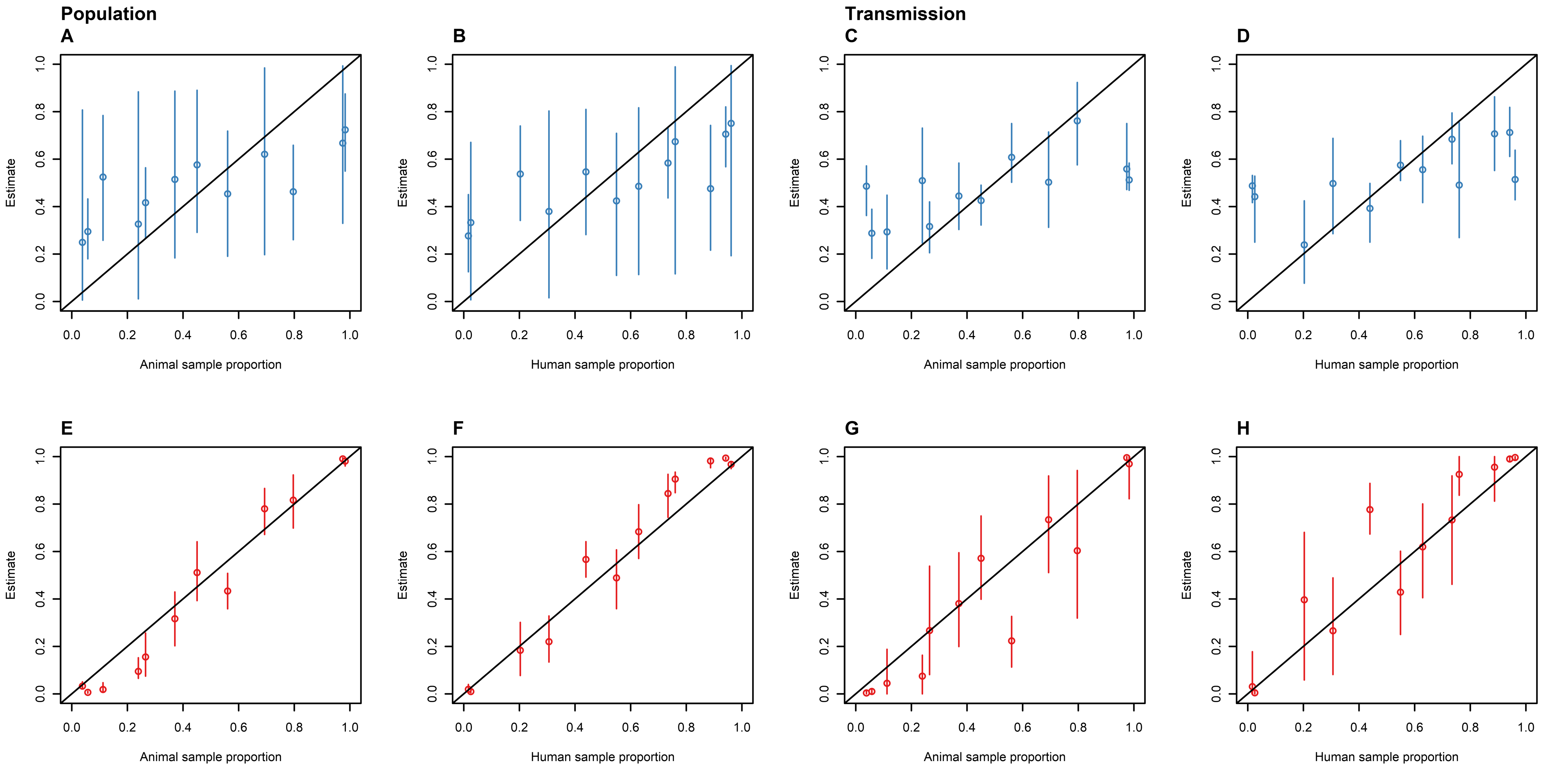

### Supplementary Figure 2

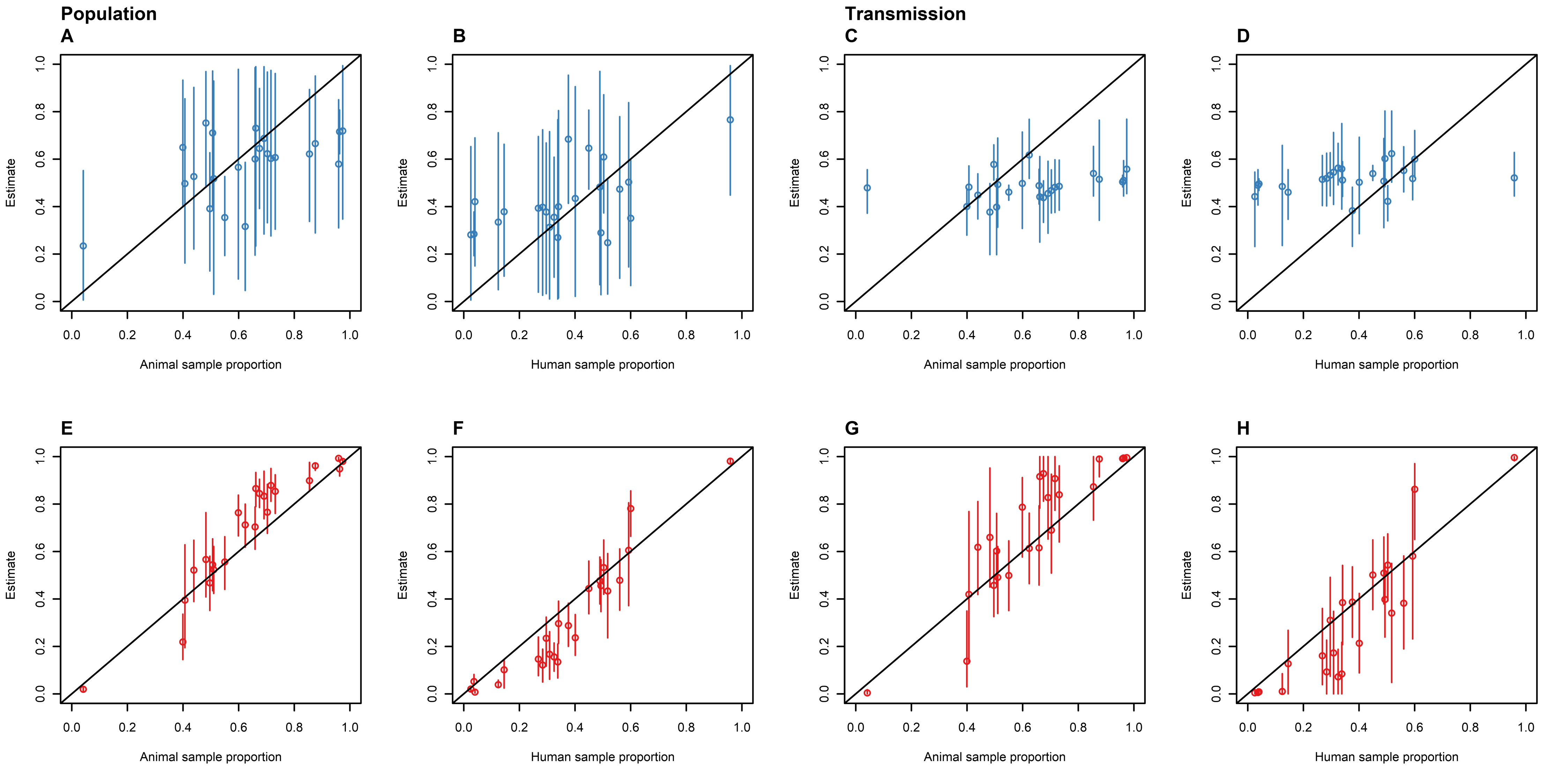

### Supplementary Figure 3

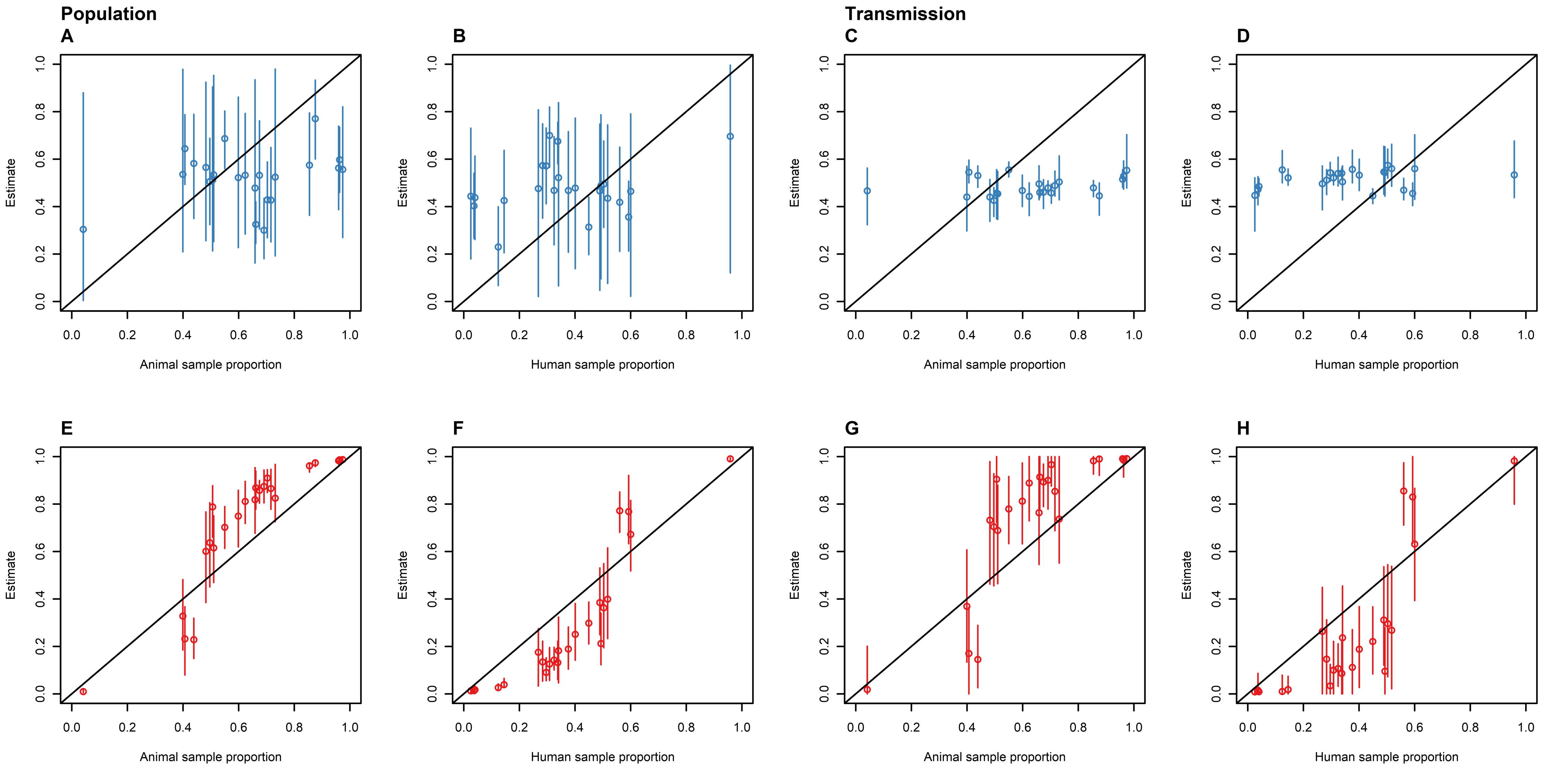

### Supplementary Figure 4

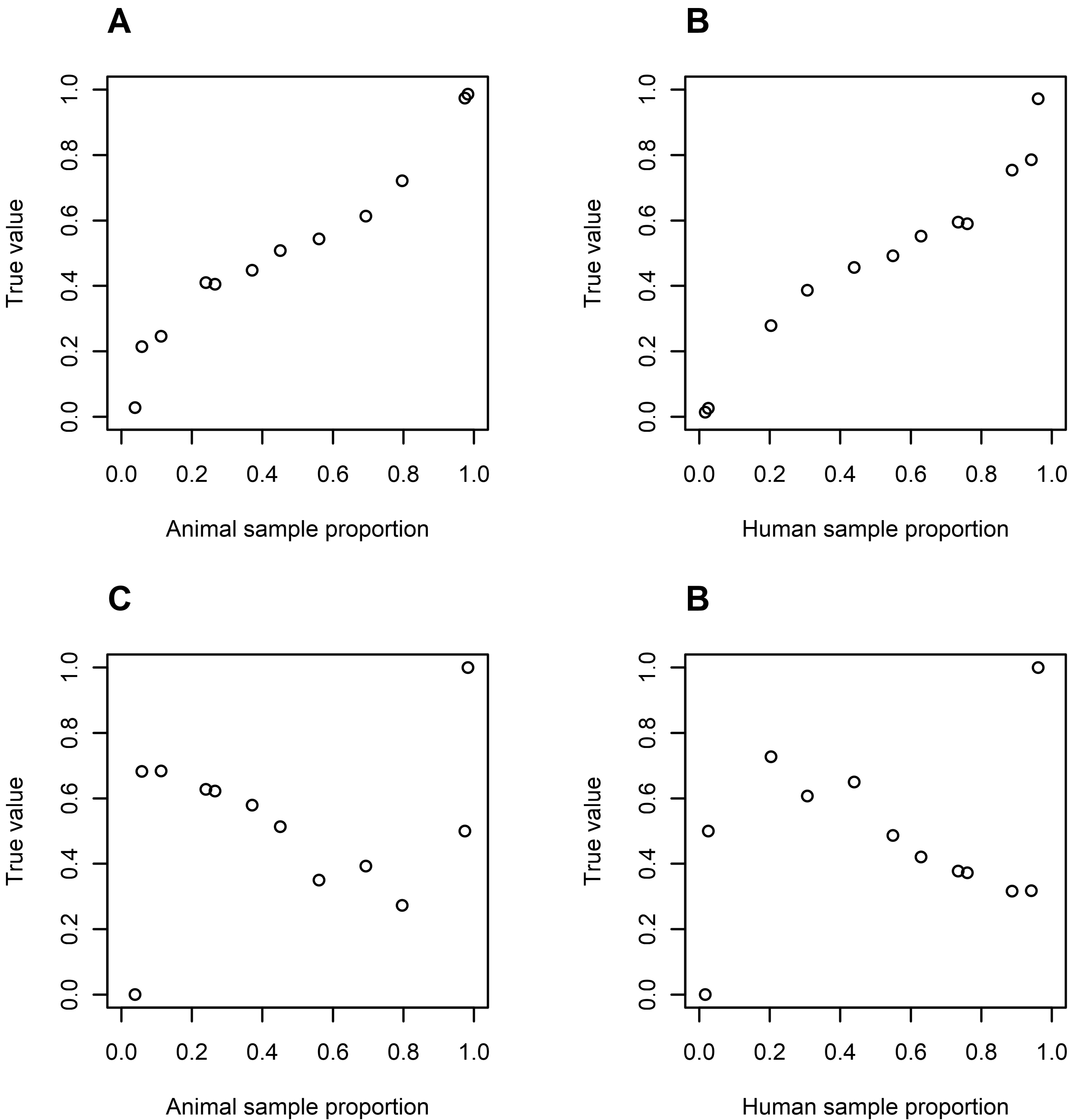

### Supplementary Figure 5

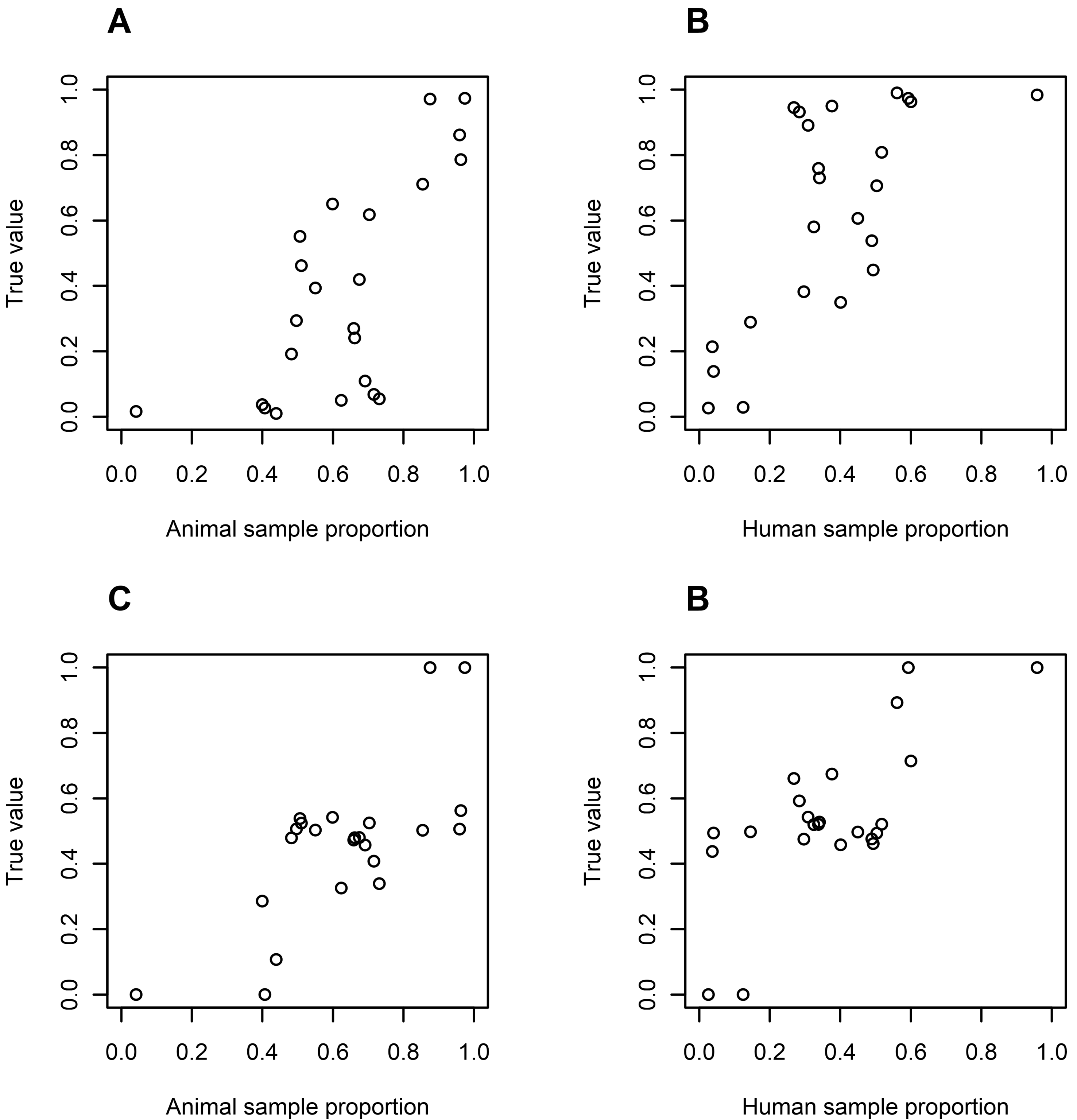

### Supplementary Figure 6

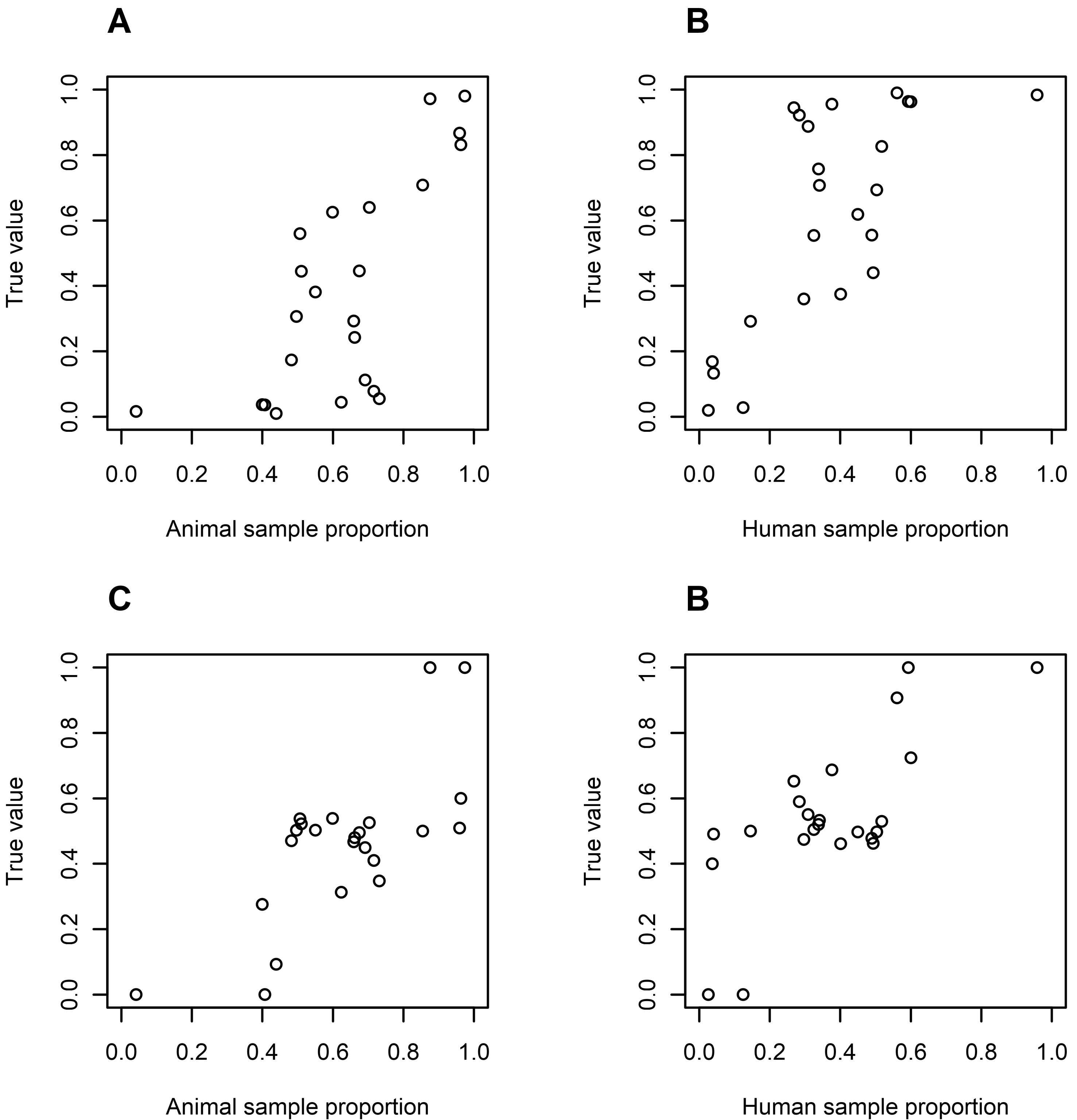
